# FACT and Ubp10 collaborate to modulate H2B deubiquitination and nucleosome dynamics

**DOI:** 10.1101/397653

**Authors:** Melesse Nune, Michael Morgan, Zaily Connell, Laura McCullough, Muhammad Jbara, Hao Sun, Ashraf Brik, Tim Formosa, Cynthia Wolberger

## Abstract

Monoubiquitination of histone H2B (H2B-Ub) plays a role in transcription and DNA replication, and is required for normal localization of the histone chaperone, FACT. In yeast, H2B-Ub is deubiquitinated by Ubp8, a subunit of SAGA, and Ubp10. Although they target the same substrate, loss of Ubp8 and Ubp10 causes different phenotypes and alters the transcription of different genes. We show that Ubp10 has poor activity on yeast nucleosomes, but that addition of FACT stimulates Ubp10 activity on nucleosomes and not on other substrates. Consistent with a role for FACT in deubiquitinating H2B *in vivo*, a FACT mutant strain shows elevated levels of H2B-Ub. Combination of FACT mutants with deletion of Ubp10, but not Ubp8, confers increased sensitivity to hydroxyurea and activates a cryptic transcription reporter, suggesting that FACT and Ubp10 may coordinate nucleosome assembly during DNA replication and transcription. Our findings reveal unexpected interplay between H2B deubiquitination and nucleosome dynamics.

## Introduction

Eukaryotic chromatin is decorated with a wide range of reversible histone post-translational modifications (PTMs) that regulate all processes that require access to DNA, including transcription, DNA replication and DNA repair (Bannister and Kouzarides, 2011; Bowman and Poirier, 2015). Actively transcribed genes in all eukaryotes are enriched in monoubiquitinated histone H2B, which plays a non-degradative role in promoting transcription (Fleming et al., 2008; Weake and Workman, 2008) but whose mechanism of action remains poorly understood. Monoubiquitin is conjugated to H2B-K123 in yeast and H2B-K120 in humans, which lie near the C-terminus of histone H2B (Robzyk et al., 2000; Weake and Workman, 2008; West and Bonner, 1980). Monoubiquitination of histone H2B is highly dynamic (Henry et al., 2003) and the cycle of ubiquitination and subsequent deubiquitination is an important checkpoint for transcription elongation (Batta et al., 2011). H2B-K123 ubiquitination is required for methylation of histone H3K4 (Chandrasekharan et al., 2010; Dover et al., 2002) and H3K79 (Ng et al., 2002), two other marks correlated with actively transcribed genes. However, H2B-Ub also plays a role in promoting transcription that is independent of cross-talk with histone H3 methylation (Tanny et al., 2007). In addition to its role in transcription, H2B-Ub plays a role in replication fork progression, nucleosome assembly during DNA replication (Ng et al., 2002) and the DNA damage response (Giannattasio et al., 2005; Moyal et al., 2011; Uckelmann and Sixma, 2017). Dysregulation of histone H2B monoubiquitination has been linked to a variety of cancers (Cole et al., 2015; Espinosa, 2008; Hahn et al., 2012).

In the yeast, *Saccharomyces cerevisiae*, histone H2B-K123 is monoubiquitinated by the E2/E3 pair, Rad6/Bre1 (Hwang et al., 2003; Robzyk et al., 2000; Wood et al., 2003), and deubiquitinated by two deubiquitinating enzymes (DUBs): Ubp8 and Ubp10 (Daniel et al., 2004; Gardner et al., 2005; Henry et al., 2003). Both of these DUBs belong to the Ubiquitin Specific Protease (USP) class of cysteine proteases, which contain a characteristic USP catalytic domain (Komander et al., 2009). Ubp10 is a monomeric enzyme whereas Ubp8 is part of a four-protein subcomplex within the SAGA complex called the DUB module, which comprises Ubp8, Sgf11, Sus1, and Sgf73 (Henry et al., 2003; Kohler et al., 2010; Samara et al., 2010b). Both yeast H2B-Ub DUBs are conserved in humans. USP36, the human homologue of Ubp10, can complement the effects on global H2B-Ub of a *ubp10* deletion in yeast (Reed et al., 2015) and USP22, the homologue of Ubp8, is a subunit of human SAGA (Zhang et al., 2008). Yeast in which both Ubp10 and Ubp8 have been deleted showed a synergistic increase in the steady-state levels of global H2B-Ub, as well as growth defects (Emre et al., 2005). While the roles of Ubp10 and Ubp8 in regulating H2B deubiquitination are well-established, their respective contributions to chromatin-mediated processes are poorly understood.

Despite their shared substrate specificity, Ubp8 and Ubp10 appear to play distinct roles *in vivo.* Several studies have shown that SAGA/Ubp8 primarily acts on H2B-Ub near promoters and transcription start sites to promote transcription initiation by RNA polymerase II (Batta et al., 2011; Daniel et al., 2004; Schulze et al., 2011). Ubp10 was first identified for its role in regulating sub-telomeric gene silencing (Emre et al., 2005; Gardner et al., 2005; Kahana and Gottschling, 1999) and is recruited to silenced chromatin (Gardner et al., 2005). However, deletion of *UBP10* alters expression of hundreds of yeast genes as well as H2B ubiquitination genome-wide (Gardner et al., 2005; Orlandi et al., 2004; Schulze et al., 2011), indicating that Ubp10 plays a global role beyond its function in subtelomeric transcriptional repression. Deletion of *UBP8* also alters transcription of several hundred genes (Gardner et al., 2005), although an analysis of the data shows little correlation between the genes whose expression is impacted by *ubp8* versus *ubp10* deletion (Figure 1). The different impacts on transcription profiles suggests that these two H2B-Ub DUBs have distinct genomic targets. However, SAGA/Ubp8 was recently shown to be involved in transcription of all RNA polymerase II genes (Baptista et al., 2017; Warfield et al., 2017) and Ubp10 has been found in association with RNA polymerase II (Mao et al., 2014), suggesting that both DUBs may at least be present at all genes. A partial resolution of this conundrum comes from a genome-wide ChIP-on-chip study of H2B-Ub in *ubp10* and *ubp8* deletion strains (Schulze et al., 2011) which shows that loss of *UBP8* results in an enrichment of H2B-Ub primarily near transcription start sites (TSS), whereas a *ubp10* deletion strain shows broader enrichment of H2B-Ub in mid-coding regions of longer transcription units. The ChIP results suggest that Ubp8 and Ubp10 are required during transcription, but at different times and in different genic locations. However, it remains unclear how each of these factors produces these distinct profiles and what roles each enzyme plays during these processes.

**Figure 1:**
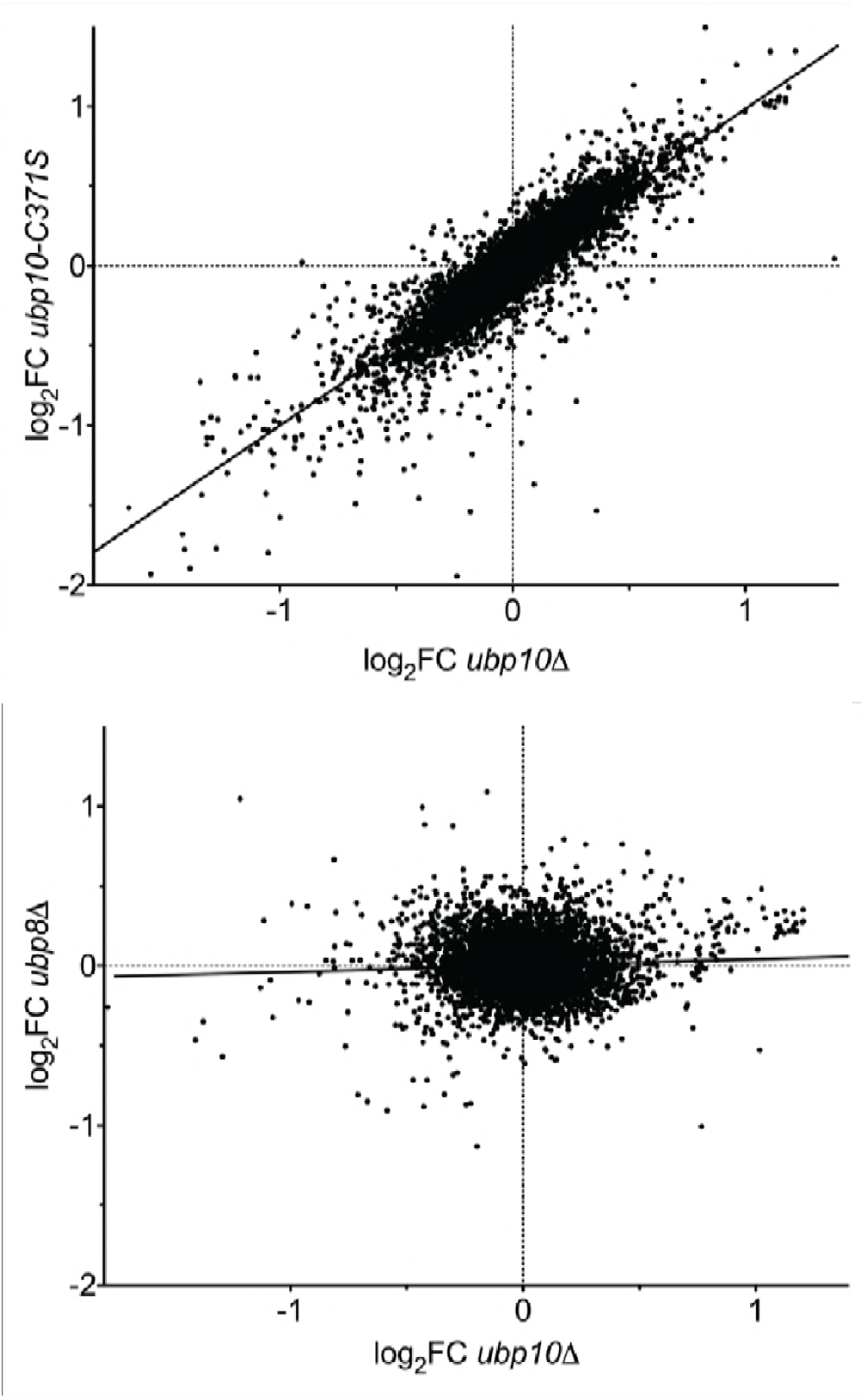
Deletion of the *UBP8* and *UBP10* genes have different effects on transcription programs. Analysis of transcription data from Gardner 2005. Scatter plots of the log_2_ fold change in transcript level vs WT (log_2_FC) are shown for (top panel) a catalytically dead allele (*ubp10-C371S*) vs a deletion (*ubp10*Δ) to demonstrate reproducibility of the array, and (bottom panel) a *ubp8*Δ strain compared with *ubp10*Δ. The two null mutants give a strong correlation (Pearson correlation coefficient r = 0.86, linear regression R^2^ = 0.74, m = 0.99), validating the reproducibility of the arrays. Deleting *UBP8* affected the transcription of different genes, resulting in poor correlation with *ubp10*Δ (r = 0.055, R^2^ = 0.0031, m = 0.039).

Ubiquitination of histone H2B has been reported to assist recruitment of the histone chaperone, FACT (Facilitates Chromatin Transcription) to active chromatin (Fleming et al., 2008). The yeast FACT complex is composed of a heterodimer of Spt16 and Pob3 that is assisted *in vitro* and *in vivo* by the DNA binding protein, Nhp6 (Brewster et al., 1998; Ruone et al., 2003a; Schlesinger and Formosa, 2000; Wittmeyer and Formosa, 1995; Wittmeyer et al., 1999). FACT evicts H2A/H2B heterodimers in front of the transcription machinery (Reinberg and Sims, 2006) and reassembles the heterodimers in the wake of RNA polymerase II to prevent cryptic transcription initiation (Fleming et al., 2008; Mason and Struhl, 2003; Pavri et al., 2006). The disruption of the H2B ubiquitination cycle or a mutation in the FACT subunit, Spt16, causes a defect in Pol II elongation (Fleming et al., 2008). In addition to roles in transcription, FACT and H2B-Ub are each also implicated in DNA replication (Formosa, 2012; Kurat et al., 2017; Trujillo and Osley, 2012). H2B-Ub at replication origins is thought to stabilize the parental nucleosomes after the passage of DNA polymerase (Trujillo and Osley, 2012). FACT and H2B-Ub play an important role in the progression of DNA replication, likely by maintaining chromatin stability and orchestrating nucleosome assembly on newly-synthesized DNA (Lin et al., 2014; Trujillo and Osley, 2012). It is clear that both FACT and H2B-Ub play a pivotal role in stabilizing and assembling nucleosomes in the wake of polymerases during replication and transcription. However, it is not known how FACT and H2B-Ub status affect one another to perform these functions.

We report here a novel role for the histone chaperone, FACT, in stimulating the H2B deubiquitination activity of Ubp10 on nucleosomes. We show that the rate of deubiquitination of yeast H2B-Ub is slower when incorporated into nucleosomes as compared to free H2A/H2B-Ub heterodimers, but that the addition of FACT reverses this block. This behavior is in marked contrast to the Ubp8/DUB module, which has robust activity on both heterodimers and intact nucleosomes and is not affected by FACT (Morgan et al., 2016). We show that a yeast strain with a FACT deficiency has elevated levels of H2B-Ub, indicating that FACT also stimulates deubiquitination of H2B *in vivo*. Deleting Ubp10, but not Ubp8, from a strain with mutated FACT conferred strong sensitivity to the DNA replication toxin, hydroxyurea (HU), and activated a cryptic transcription reporter. Our findings suggest that the differential effects of Ubp10 and Ubp8 on the distribution of H2B-Ub result from a global role for Ubp10 and FACT versus a local role of Ubp8/SAGA at promoters and transcription start site. These observations have important implications for the way in which cycles of H2B ubiquitination and deubiquitination regulate nucleosome dynamics during transcription and DNA replication.

## Results

### Ubp10 preferentially deubiquitinates free yeast H2A/H2B-Ub relative to nucleosomes

During transcription, nucleosomes are at least partially disassembled in order to enable RNA polymerase to access the DNA template and are then reassembled in the wake of the transcribing polymerase. It is not known when during this process ubiquitin is conjugated to histone H2B and when it is removed by either Ubp8/SAGA or Ubp10. Since histone H2A/H2B heterodimers can be ejected and re-inserted during the dynamic nucleosome disassembly and reassembly that accompanies passage of RNA polymerase, it is formally possible that H2B is deubiquitinated when it is in an intact nucleosome, after ejection to the free H2A/H2B-Ub dimer form, or when the nucleosome is in an intermediate state of disassembly or assembly. We previously reported that the Ubp8/SAGA DUB module deubiquitinates H2B in the context of both the nucleosome and the free H2A/H2B-Ub heterodimers, with a modest preference for nucleosomes (Morgan et al., 2016). Those results suggested that Ubp8/SAGA could deubiquitinate H2B at any point during transcription.

Since Ubp10 has been reported to associate with RNA polymerase II (Mao et al., 2014) and to deubiquitinate H2B in gene bodies (Schulze et al., 2011), we asked whether this monomeric DUB discriminates between H2B-ubiquitinated nucleosomes and ubiquitinated H2A/H2B-Ub heterodimers. Using an intein-based semisynthetic approach, we generated ubiquitinated yeast histone H2B in which the C-terminus of ubiquitin was linked to H2B-K123 via a native isopeptide linkage (Jbara et al., 2018; Maity et al., 2016). This H2B-Ub was used to reconstitute nucleosomes and H2A/H2B-Ub heterodimers. Remarkably, Ubp10 cleaved ubiquitin from H2A/H2B-Ub heterodimers at least 100-fold faster than from nucleosomes containing H2B-Ub (Figure 2A). Under the conditions tested, the majority of H2A/H2B-Ub was consumed in less than 5 minutes while almost all of the NCP-Ub remain uncleaved after 60 minutes. Similar behavior was recently observed in experiments using human histones containing a cleavable analogue of a native isopeptide linkage and a GST-Ubp10 fusion (Zukowski et al., 2018). Taken together, our results indicate that Ubp10 discriminates between freestanding histone heterodimers and those in nucleosomes (Figure 2A) whereas Ubp8/SAGA does not (Morgan et al., 2016).

**Figure 2:**
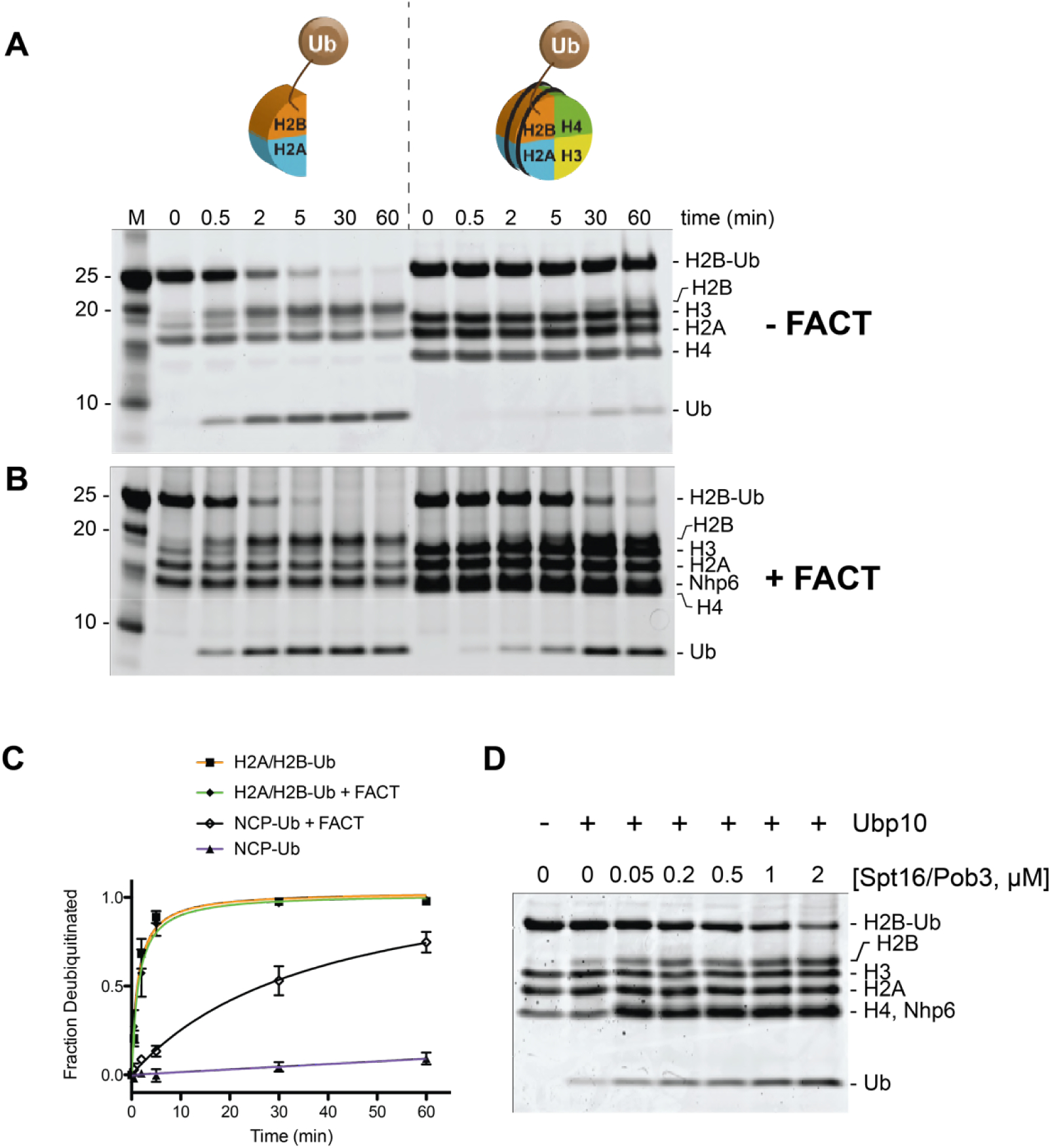
Ubp10 preferentially deubiquitinates H2A/H2B-Ub over nucleosomal (NCP) H2B-Ub. A) Ubp10 activity on H2A/H2B-Ub and NCP-Ub in the absence of FACT. 1 µM NCP-Ub and 2 µM H2A/H2B-Ub were incubated with 5 nM Ubp10 and time points were taken by quenching with SDS sample buffer. B) FACT stimulates Ubp10 activity on NCP-Ub. Ubp10 activity was measured as in A, but in the presence of FACT subunits, 2 µM Spt16/Pob3-WT and 2 µM Nhp6. C) Plot showing the fraction of total substrate consumed over time. The plot was generated by averaging the relative intensity of H2B-Ub bands as compared with that of uncleaved H2B-Ub at t=0 from 3 independent experiments (mean normalized band intensity and standard deviation shown). D) Increasing the concentration of FACT increases the activity of Ubp10. 1 µM NCP-Ub was incubated with the indicated concentrations of Spt16/Pob3-WT and 2 µM Nhp6 in the presence of 5 nM Ubp10. Each reaction was quenched at 60 minutes.

### FACT stimulates Ubp10 DUB activity on nucleosomes

In cells, core histones that are not incorporated into nucleosomes are usually bound by histone chaperones, which bind to H2A/H2B heterodimers or H3/H4 heterodimers or heterotetramers (Elsasser and D’Arcy, 2013). The histone chaperone, FACT, binds to H2A/H2B heterodimers and facilitates heterodimer eviction and exchange, as well as nucleosome reassembly (Fleming et al., 2008; Mason and Struhl, 2003; Orphanides et al., 1998; Orphanides et al., 1999; Saunders et al., 2003). In light of the reported functional interaction between H2B-Ub and FACT (Fleming et al., 2008; Pavri et al., 2006) and the role of FACT in binding both H2A/H2B heterodimers and intact nucleosomes, we asked whether Ubp10 can remove ubiquitin from ubiquitinated yeast H2A/H2B heterodimers or nucleosomes when they are bound to FACT. We unexpectedly found that FACT dramatically increases the rate at which Ubp10 cleaves H2B-Ub in nucleosomes (Figure 2B-C). Essentially all of the H2B-Ub in the nucleosomal sample was cleaved in under 60 minutes in the presence of FACT (Figure 2B-C), whereas less than 10% was consumed during the same time period in the absence of FACT (Figure 2A-C). By contrast, the addition of FACT had no effect on the rate at which Ubp10 deubiquitinated H2B in H2A/H2B-Ub heterodimers (Figure 2A-B). We verified that purified FACT on its own has no DUB activity against the ubiquitinated nucleosome (Figure2 – Figure supplement 1). To further confirm the dependence of Ubp10 DUB activity on the presence of FACT, we assayed Ubp10 deubiquitination activity on nucleosomes at a fixed time point in the presence of increasing concentrations of FACT. As shown in Figure 2D, the amount of nucleosomal H2B-Ub cleaved increases as a function of increasing FACT concentration. Notably, the dose response for FACT in this assay closely matches the affinity of FACT for nucleosomes (Ruone et al., 2003b; Winkler et al., 2011). These results show that FACT stimulates Ubp10 DUB activity, and that this stimulatory effect is specific to nucleosomal H2B-Ub substrates.

A possible explanation for the observed stimulatory effect of FACT is that it alters nucleosomal structure, making it a better substrate for Ubp10. Previous studies have shown that FACT binding can destabilize the canonical nucleosomes, disrupting the octamer/DNA contacts, which could result in displacement of H2A/H2B heterodimers (Belotserkovskaya et al., 2003; Chen et al., 2018; McCullough et al., 2011), thereby providing better substrates for Ubp10 (Figure 2). During nucleosome reorganization induced by FACT, surfaces of H2A/H2B heterodimers that are buried in the context of the nucleosome become more accessible (Kemble et al., 2015) even while the components remained tethered together (Wang et al., 2018; Xin et al., 2009a; Yang et al., 2018), which could enhance accessibility of H2B-Ub for deubiquitination by Ubp10 even without dimer eviction. We therefore tested the simple model in which binding of DNA to H2A/H2B heterodimers creates a barrier to Ubp10 deubiquitination of H2B-Ub. However, the activity of Ubp10 on H2A/H2B-Ub heterodimers was similar in the presence and absence of 601 DNA (Figure 2 – Figure supplement 2). This experiment does not rule out steric hindrance by DNA in the context of the full histone octamer, but leaves open the possibility that reorganization of the nucleosome by FACT exposes surfaces more favorable to Ubp10 docking on H2B-Ub.

The finding that FACT stimulates Ubp10 to deubiquitinate nucleosomes stands in stark contrast to the Ubp8/SAGA DUB module. We previously found (Morgan et al., 2016) that the abillity of the Ubp8/DUB module to deubiquitinate nucleosomes is not affected by the addition of FACT. SAGA/Ubp8 therefore can access H2B-K123Ub in the context of the nucleosome, but Ubp10 is unable to deubiquitinate nucleosomal H2B-Ub without the assistance of FACT.

### FACT stimulation does not correlate with Ubp10 nucleosome-binding activity

Ubp10 contains an unstructured region rich in Asp/Glu that is N-terminal to the catalytic USP domain (residues 362 - 733) (Reed et al., 2015) (Figure 3A). The N-terminal unstructured region contains residues that interact with the Sir3/Sir4 silencing proteins and recruit Ubp10 to subtelomeric regions (Emre et al., 2005; Gardner et al., 2005; Zukowski et al., 2018). However, it is not known how Ubp10 is recruited to ubiquitinated nucleosomes elsewhere in the genome. We therefore asked whether Ubp10 alone can bind to nucleosomes in which ubiquitin is linked to H2B-K123 via a non-hydrolyzable linker (Morgan et al., 2016). We detected binding of Ubp10 to ubiquitinated nucleosomes in an electrophoretic mobility shift assay (EMSA), with half-maximal binding observed at approximately 0.4 µM Ubp10 (Figure 3B). Ubp10 bound with similar apparent affinity to unmodified nucleosomes (Figure 3 – Figure supplement 1), indicating that interactions between the ubiquitin and the Ubp10 catalytic domain do not play a significant role in the observed binding. Deletion of the N-terminal 156 residues had little effect on the affinity of Ubp10 for ubiquitinated nucleosomes (Figure 3B). However, a further deletion of the N-terminal 199 residues, Ubp10-(200-792), reduced binding substantially, and no binding was detected with Ubp10 residues 250-792 (Figure 3B). These results show that the Asp/Glu rich region (residues 157-250) of Ubp10 is important for the observed binding to nucleosomes.

**Figure 3:**
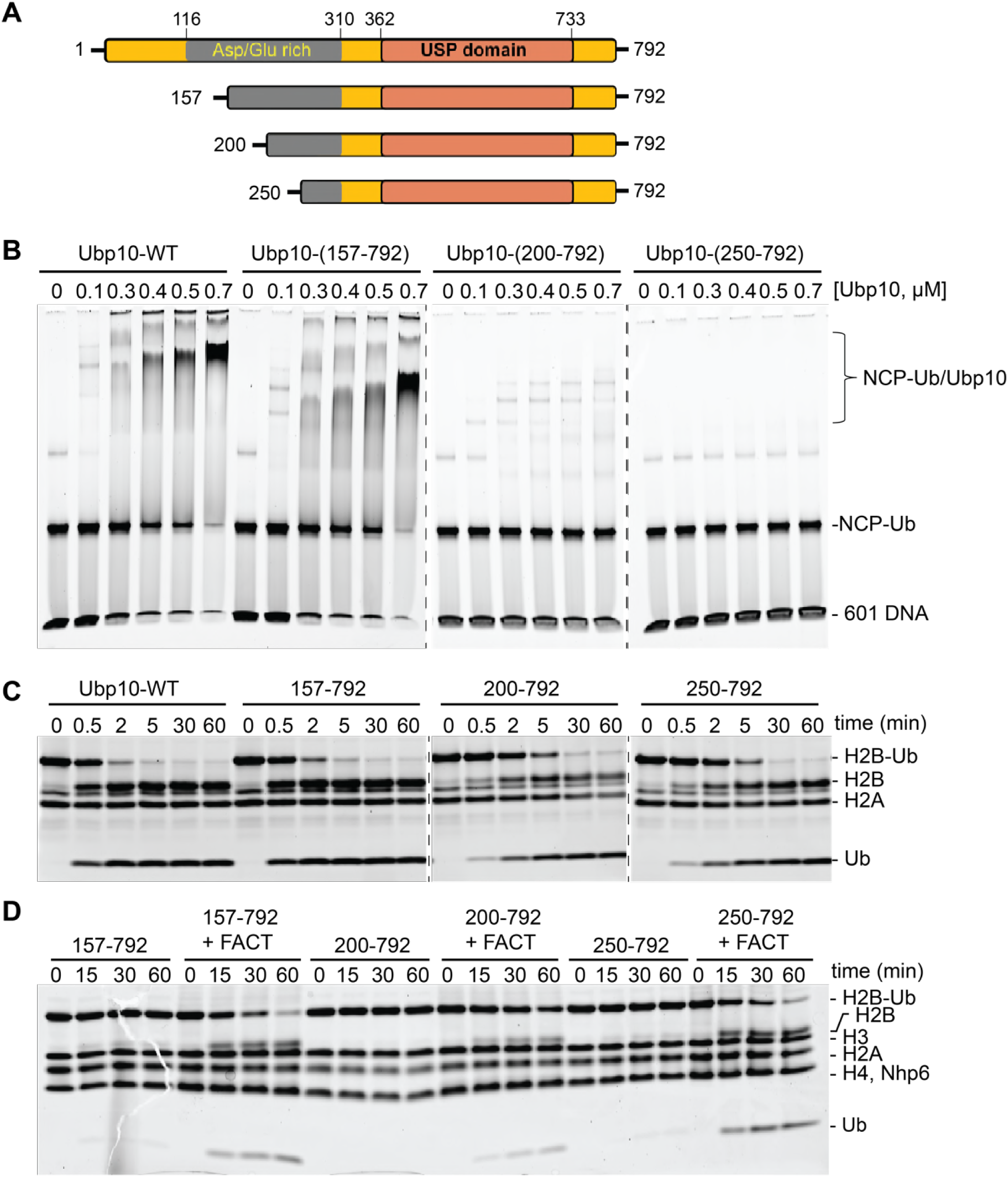
FACT stimulates Ubp10 truncations defective in binding to nucleosomes. A) Schematic of Ubp10 and truncations used showing the catalytic domain (USP) and an N-terminal region rich in aspartic acid and glutamic acid repeats. B) Native gel showing complexes formed between Ubp10 constructs and ubiquitinated nucleosomes. C) Ubp10 activity was measured as in Figure 1; time points were taken after incubating 2 µM H2A/H2B-Ub dimers with 5 nM Ubp10 fragments. D) NCP-Ub cleavage activity monitored in the presence and absence of FACT and several Ubp10 constructs.

To test whether Ubp10 domains that are required for nucleosome binding are also required for DUB activity as well as for stimulation by FACT, we tested the DUB activity of the Ubp10 N-terminal truncations. All three N-terminal truncation mutants were active on H2A/H2B-Ub heterodimers, although the Δ200 and Δ250 truncations had slightly lower activity than intact Ubp10 (Figure 3C). Similar to the full length protein, all three truncations displayed weak activity on a nucleosomal substrate in the absence of FACT and enhanced DUB activity on nucleosomes in the presence of FACT (Figures 3D and 2A). Therefore, the truncations that decreased Ubp10 affinity for nucleosomes (Ubp10 200-792 and Ubp10 250-792) were still stimulated by FACT. We also tested the hypothesis that the presence of FACT might enhance binding of Ubp10 to nucleosomes, which could provide a mechanism by which FACT enhances cleavage of nucleosomal H2B-Ub. Gel mobility shift assays for binding to ubiquitinated nucleosomes, however, did not show enhanced affinity of Ubp10 for ubiquitinated nucleosomes in the presence of FACT (Figure 3 – Figure supplement 2). These results suggest that the ability of FACT to stimulate Ubp10 is not coupled to the intrinsic ability of Ubp10 to bind nucleosomes.

### A FACT mutant strain has elevated levels of H2B-Ub

Our *in vitro* assays show that the ability of Ubp10 to deubiquitinate nucleosomes is greatly enhanced in the presence of the histone chaperone FACT (Figure 2). If FACT activity also stimulates Ubp10 activity *in vivo*, a defect in FACT activity should phenocopy the effects of a Ubp10 deletion. We compared the relative ratio of monoubiquitinated H2B to unmodified H2B in a *pob3-L78R* yeast strain, which contains a destabilizing mutation in the Pob3 subunit of FACT that reduces FACT levels by about 10-fold under permissive growth conditions (Schlesinger and Formosa, 2000; VanDemark et al., 2008). This mutant has been shown to have defects in both transcription and DNA replication (Schlesinger and Formosa, 2000). As shown in Figure 4, the *pob3-L78R* strain had an elevated level of H2B-Ub that is comparable to that in a *ubp10* deletion strain when normalized for total H2B, which varies due to changes in cell size and thus relative contribution of nuclear proteins to the total. This increase in H2B-Ub in a FACT mutant is consistent with a role for FACT in deubiquinating H2B-Ub *in vivo*. A strain lacking Ubp10 and also carrying the FACT defect had roughly the same increased level of H2B-Ub as *the ubp10* mutant and the same ratio of increase as the *pob3-L78R* mutant (Figure 4B), consistent with the interpretation that FACT and Ubp10 cooperate to deubiquitinate H2B in the same pathway. Together, these results show that FACT activity can contribute to H2B deubiquitination *in vivo*, supporting the physiological relevance of the *in vitro* data showing that FACT is required to stimulate Ubp10 activity on nucleosomes (Figure 2D).

**Figure 4:**
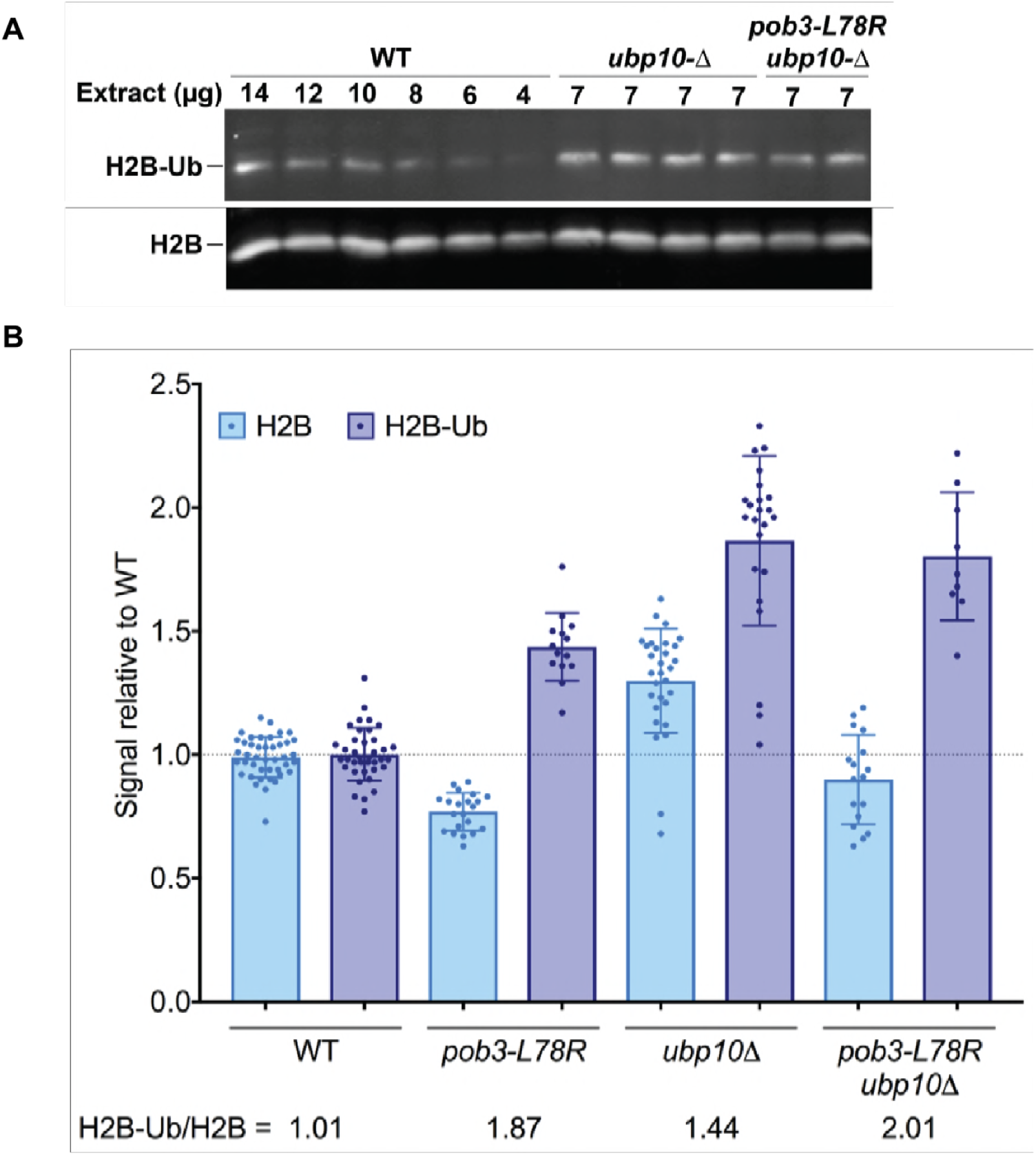
H2B-Ub levels are elevated in a FACT mutant strain. A) Representative Western blot analysis of TCA extracts from *WT*, *ubp10Δ*, and *ubp10Δ pob3-L78R* strains probed with antibodies against H2B-Ub and re-probed with antibodies against H2B. B) Relative steady-state levels of H2B-Ub for *WT*, *pob3-L78R, ubp10Δ, and ubp10Δ/pob3-L78R* strains (Table 1). The average and standard deviation from multiple biological replicates is shown. The numbers at the bottom indicate the relative H2B-Ub increase normalized to unmodified H2B (H2B-Ub/H2B) within each individual gel.

### FACT and Ubp10 cooperate during DNA replication

In addition to their roles in transcription (Batta et al., 2011; Fleming et al., 2008; Pavri et al., 2006; Reinberg and Sims, 2006; Weake and Workman, 2008), both FACT and H2B monoubiquitination have been implicated in assembling and stabilizing nucleosomes during DNA replication (Jasencakova and Groth, 2010; Lin et al., 2014; Trujillo and Osley, 2012). FACT has been proposed to play an important role in DNA replication by assisting in initiation, DNA unwinding, histone eviction, and chromatin reassembly (Lin et al., 2014; Ransom et al., 2010; Schlesinger and Formosa, 2000; Trujillo and Osley, 2012). H2B monoubiquitination near origins of replication supports replisome stability, fork progression and checkpoint pathways (Lin et al., 2014; Trujillo and Osley, 2012). Insights into the role of H2B-Ub in DNA replication come from studying the effects of deleting the H2B E3 ligase, Bre1 (Lin et al., 2014; Trujillo and Osley, 2012), or mutants expressing H2B with a K123R substitution, which cannot be ubiquitinated (Lin et al., 2014; Trujillo and Osley, 2012). The role of H2B deubiquitinating enzymes in DNA replication, however, has not been explored.

**Table 1.**
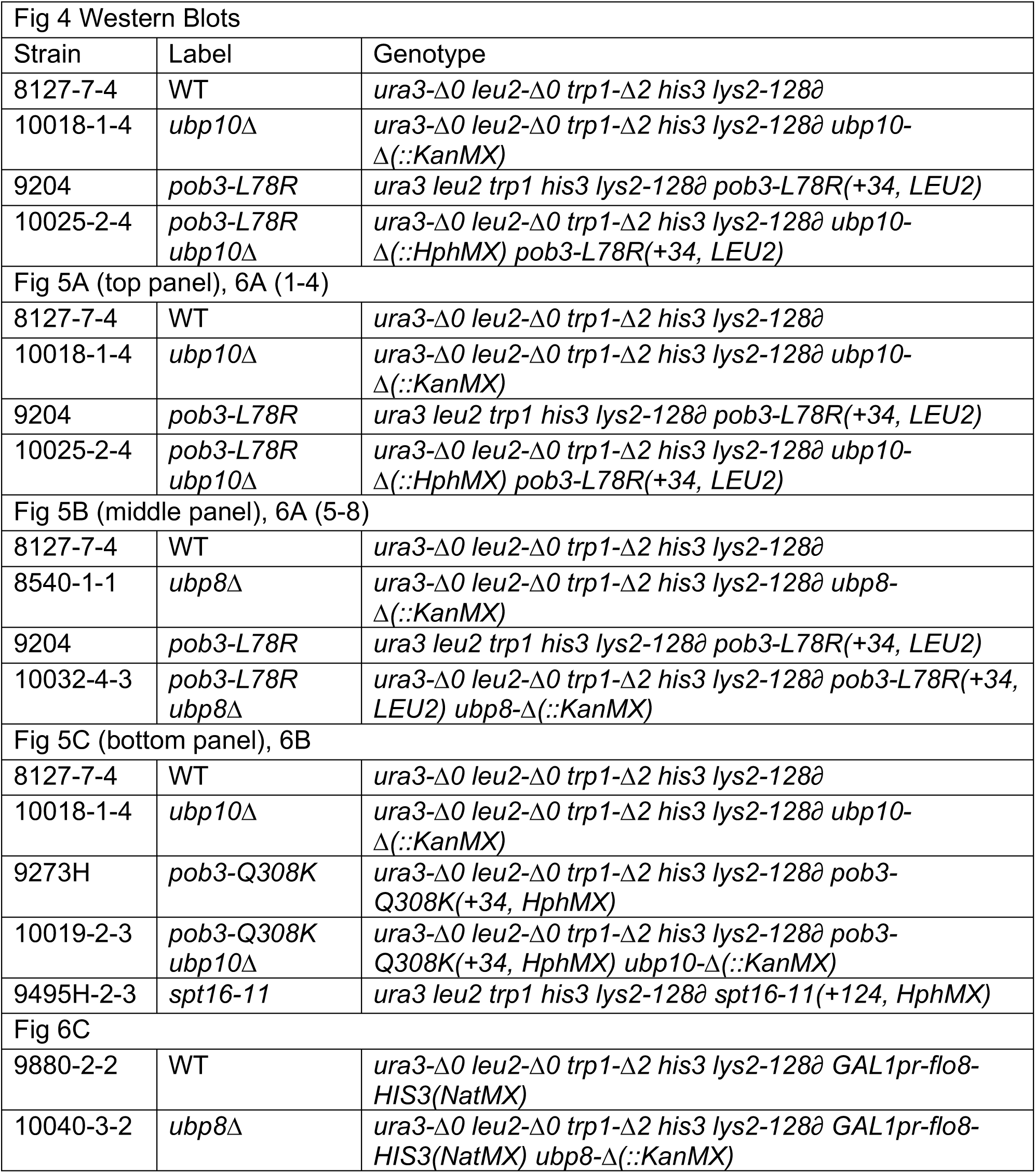
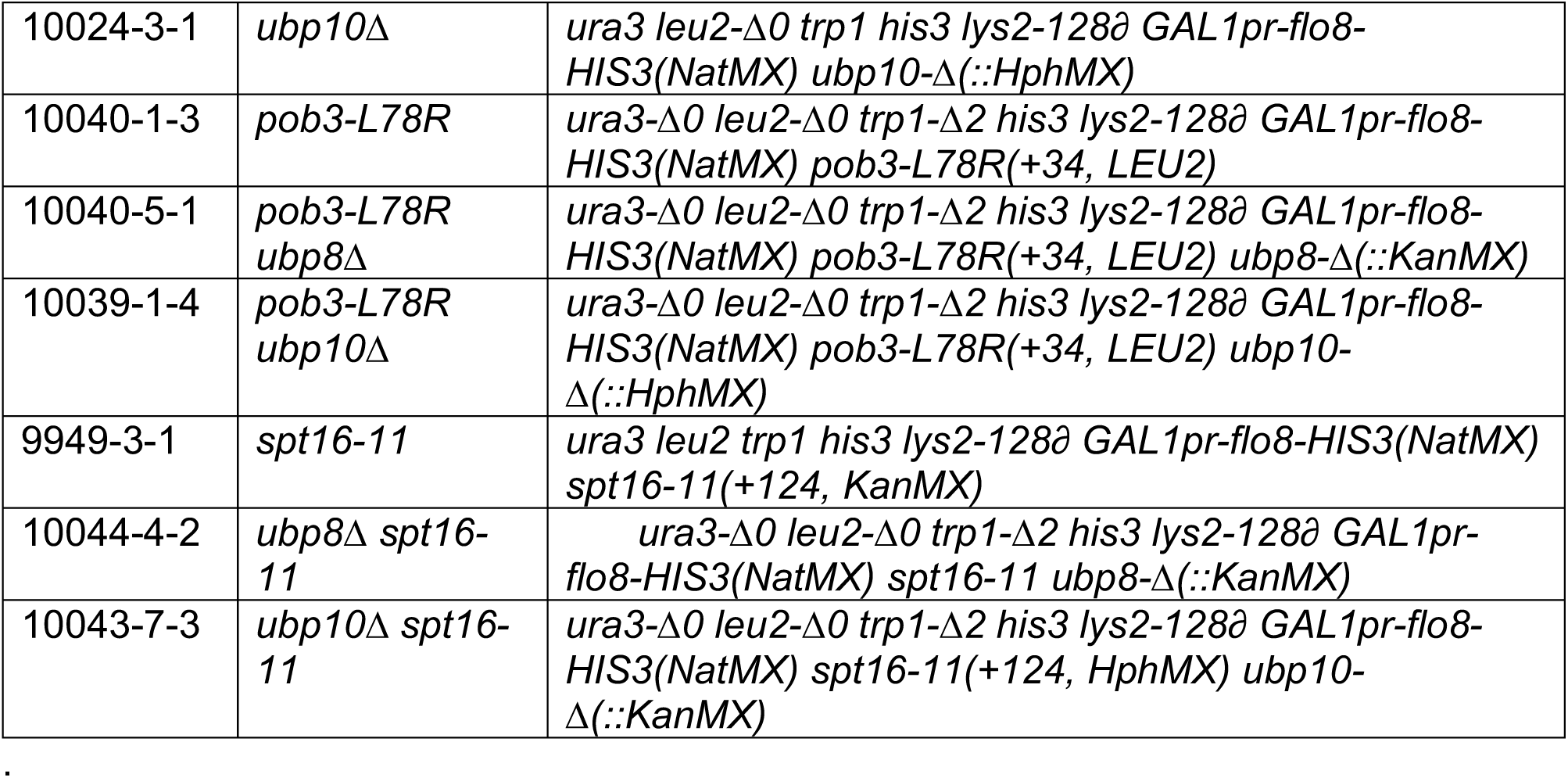
Yeast Strains used. All strains are congenic with the A364a background and are *MATa.* Standard methods were used to introduce the mutations shown into diploid strains, then haploids were derived and crossed to obtain the combinations listed, ensuring that all strains with the same genotype displayed the phenotypes observed.

In light of our finding that FACT stimulates Ubp10 activity on nucleosomes *in vitro* and is required to maintain wild type levels of H2B ubiquitination *in vivo*, we asked whether FACT functionally interacts with Ubp10 during DNA replication. We tested the sensitivity of Ubp10 deletions and FACT mutants to hydroxyurea, which depletes cellular dNTPs and induces replication fork stalling (Singh and Xu, 2016). The *pob3-L78R* FACT mutant has previously been shown to display a severe growth defect at 30° (temperature sensitivity) but is viable at 25°C (Schlesinger and Formosa, 2000), while the *Δubp10* deletion strain has a mild growth defect and is not temperature sensitive (Gardner et al., 2005). Growth of the *Δubp10/pob3-L78R* double mutant at 25°C was comparable to that of *pob3-L78R* alone (Figure 5A). Neither the *Δubp10* nor the *pob3-L78R* FACT mutant was sensitive to 90 mM hydroxyurea (Figure 5A). However, the double *Δubp10/pob3-L78R* mutant displayed a significant growth defect under these or milder conditions, with a synthetic defect observed even at levels of HU as low as 6 mM (Figure 5A). To test whether this synthetic growth defect in the presence of HU is unique to the *pob3-L78R* FACT mutant, we also examined the genetic interaction between a *ubp10* deletion and two other FACT mutants, *spt16-11* and *pob3-Q308K*. As shown in Figure 5C, both the *spt16-11* and *pob3-Q308K* mutants displayed synthetic growth defects in the presence of HU when combined with a *ubp10* deletion.

**Figure 5:**
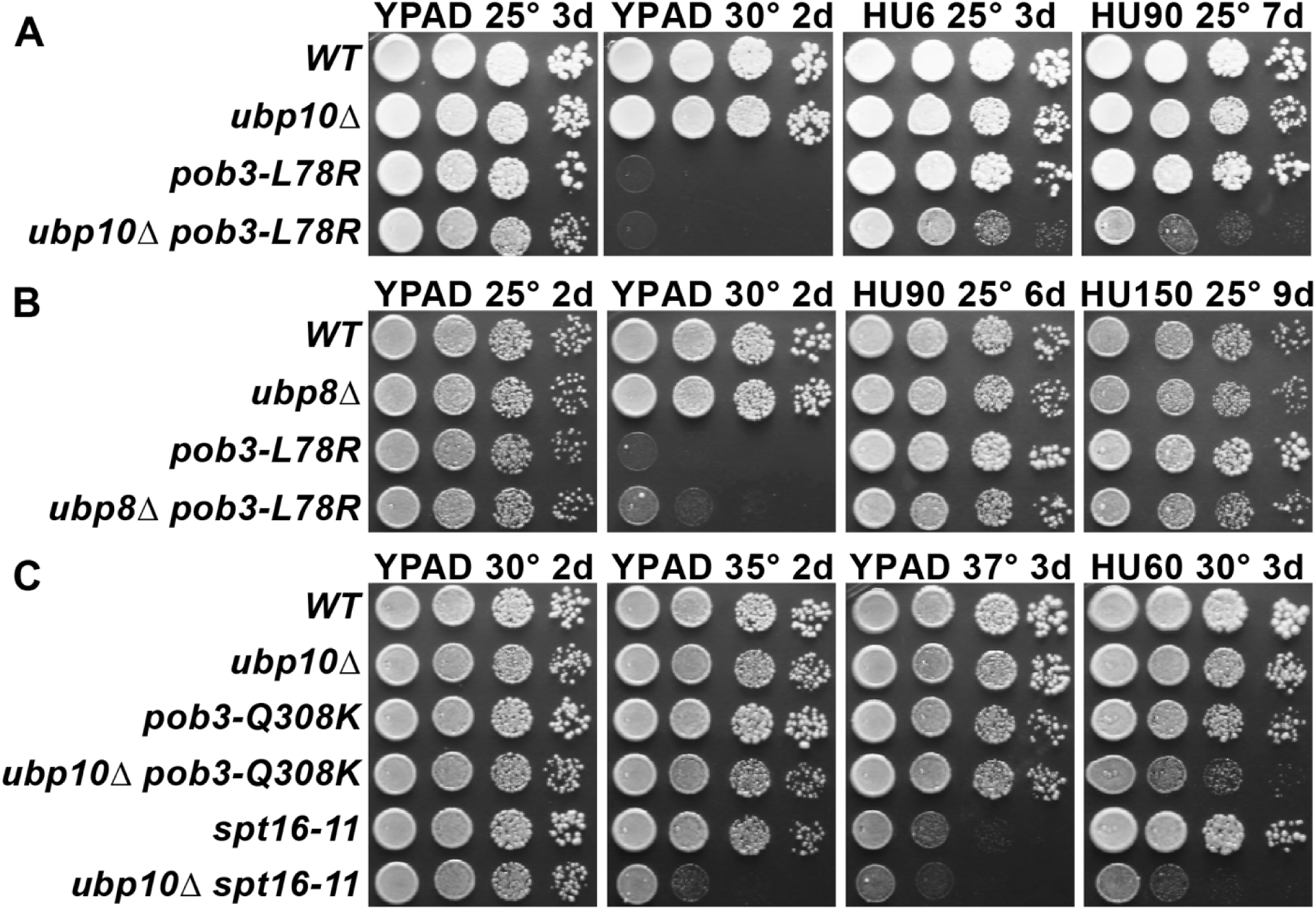
Combining *ubp10*Δ and FACT alleles causes sensitivity to HU. Strains indicated (Table 1) were grown to saturation, then 10-fold serial dilutions were spotted on rich medium (YPAD) with and without the indicated concentrations of hydroxyurea (HU, mM). Plates were incubated at the temperature indicated for the time shown (days). Combining *pob3-L78R* with *ubp10*Δ caused HU sensitivity (A), but combining it with *ubp8*Δ did not (B). Combining *ubp10*Δ with other alleles of FACT also caused synthetic defects on HU (C).

Ubp8/SAGA, in contrast to Ubp10, does not depend upon FACT to deubiquitinate nucleosomes *in vitro* (Morgan et al., 2016). We therefore predicted that *ubp8* and FACT should not have the same genetic interaction as *ubp10* and FACT. Indeed, a *Δubp8/pob3-L78R* double mutant did not show any synthetic growth defects in the presence of hydroxyurea, even at concentrations as high as 150 mM (Figure 5C). These observations point to a specific cooperative function of Ubp10 and FACT in DNA replication that cannot be performed by Ubp8.

### FACT and Ubp10 cooperate to suppress cryptic transcription

Both FACT and H2B ubiquitination are needed to maintain wild type levels of nucleosome occupancy (Fleming et al., 2008; Jamai et al., 2009). Defects in nucleosome occupancy can give rise to altered transcription patterns and activation of cryptic transcription initiation in gene coding regions (Fleming et al., 2008; Kaplan et al., 2003). Mutations in FACT cause expression of the *lys2-128∂* Spt reporter (<u>S</u>u<u>p</u>pression of <u>T</u>y1 insertion) (Brewster et al., 1998; Malone et al., 1991; Schlesinger and Formosa, 2000; Simchen et al., 1984), which reveals failure to establish normal chromatin-mediated repression of this promoter (Cheung et al., 2008; Kaplan et al., 2003). Wild type strains with this reporter have normal chromatin and do not grow on medium lacking lysine, but all three FACT mutants tested here express the reporter and grow (called the Spt– phenotype; Figure 6). This readout of transcription initiation resulting from poor quality chromatin was not affected by the loss of Ubp10 (Figure 6A, B; the growth defect caused by combining *ubp10Δ* with FACT mutations lead to a similar decrease in growth on -lys and complete media for double mutants). Combining a *ubp8Δ* allele with either FACT mutation, by contrast, had no effect. We then tested the effect of *ubp10Δ* on expression of a cryptic promoter reporter, which supports growh in the absence of histidine if galactose-induced transcription of the reporter activates a cryptic promoter within the *FLO8* gene (Cheung et al., 2008) (Figure 6D). While neither a *ubp10Δ* nor *spt16-11* mutant activated the reporter gene, the double *Δubp10/spt16-11* mutant displayed significant levels of cryptic promoter activation (compare Gal complete and Gal-his in Figure 6) whereas the double *Δubp8/spt16-11* mutant did not. We were not able to see an effect of combining a *ubp10Δ* with the *pob3-L78R* allele because the *pob3-L78R* mutation on its own robustly activates the cryptic transcription reporter and enables near wild-type levels of growth (Figure 6). Taken together, these results indicate that Ubp10, but not Ubp8, acts in combination with FACT to maintain normal chromatin organization in the wake of RNA polymerase II transcription.

**Figure 6:**
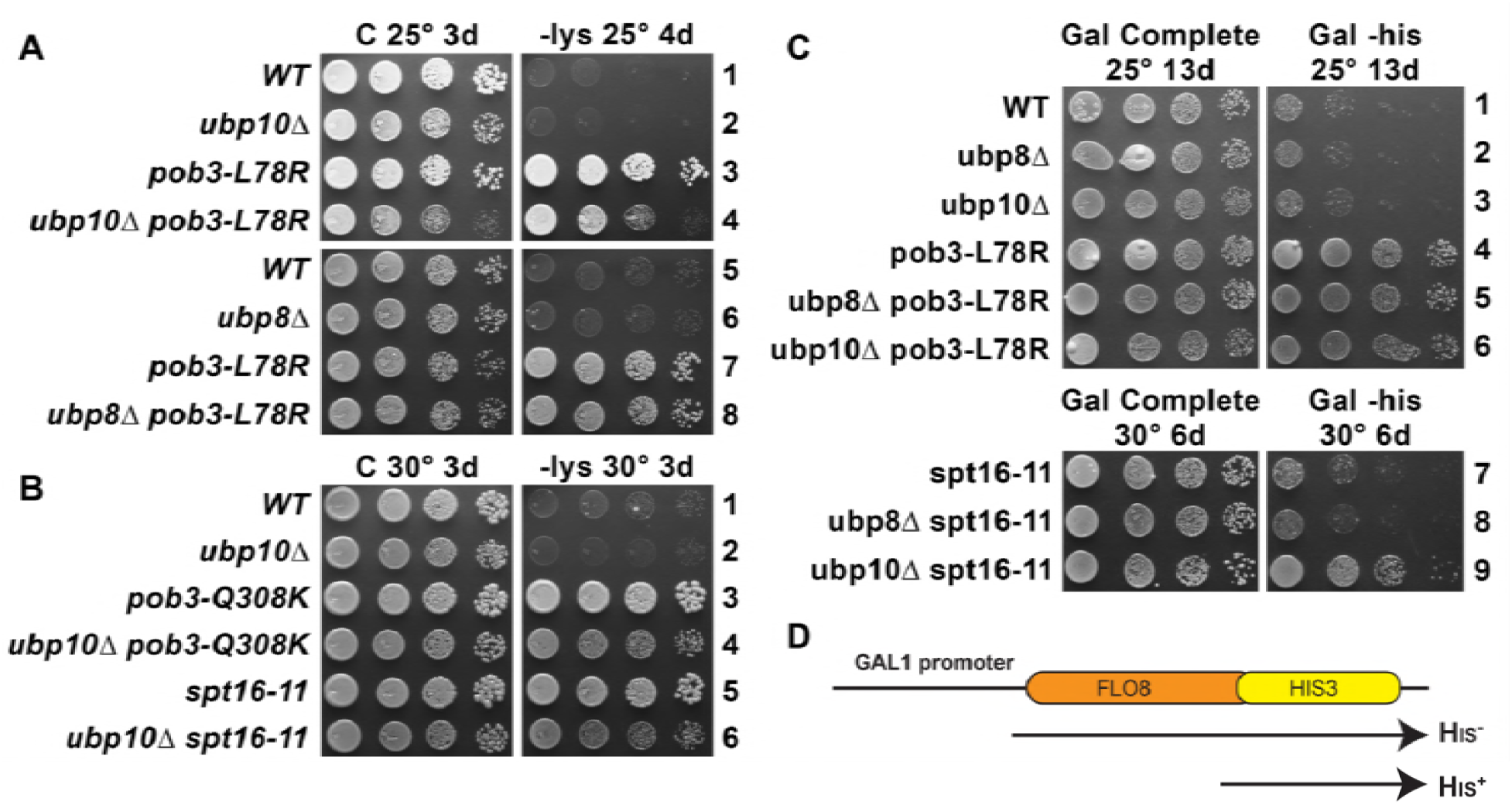
Combined effect of FACT mutants and *ubp10*Δ or *ubp8*Δ deletion on Spt-phenotype and cryptic promoter activation. A,B) *ubp10*Δ did not affect the Spt– phenotype of FACT mutants. Dilutions of the same strains shown in Figure 5 were plated on synthetic medium with (C, complete) or without lysine (-lys). These strains with the *lys2-128∂* are auxotrophic for lysine, but defects in chromatin integrity allow expression of this allele revealed by growth onlys (the Spt– phenotype; see Winston 2008). FACT mutants displayed this phenotype but this was not affected by *ubp8*Δ or *ubp10*Δ (A, B, and not shown). C) Activation of a cryptic transcription reporter in a *ubp10*Δ/*spt16-11* mutant strain reveals a defect in restoring chromatin in the wake of RNA Pol II passage. Strains with an out-of-frame fusion of *HIS3* to a site downstream of a cryptic promoter in the *FLO8* gene (panel D, adapted from Cheung et al. 2008) are auxotrophic for histidine when the *GAL1* promoter driving transcription of this reporter is repressed on glucose (not shown) but can grow without histidine on synthetic medium containing galactose (Gal -his). Strains with the *pob3-L78R* mutation have this phenotype, indicating activation of the cryptic promoter, masking any potential effects of *ubp8*Δ or *ubp10*Δ. The *spt16-11* allele alone did not activate this reporter but did when combined with *ubp10*Δ.

## Discussion

We have discovered a previously uncharacterized synergy between the H2B deubiquitinating enzyme, Ubp10, and the histone chaperone, FACT, that suggests a unifying model for the role of H2B ubiquitination and deubiquitination in nucleosome dynamics. The role of H2B monoubiquitination in recruiting FACT to nucleosomes and the importance of both FACT and H2B-Ub in transcription (Batta et al., 2011; Fleming et al., 2008; Weake and Workman, 2008) and in DNA replication (Lin et al., 2014; Trujillo and Osley, 2012), are well-established. However, the interplay between nucleosome dynamics, H2B deubiquitination and histone chaperones had not been explored. In this study, we report a role for the histone chaperone, FACT, in deubiquitinating H2B-Ub and show that Ubp10 depends on FACT to efficiently cleave H2B-Ub from nucleosomes (Figure 2). This behavior is unlike Ubp8/SAGA, which does not require FACT to deubiquitinate nucleosomes (Morgan et al., 2016). Consistent with our *in vitro* results, we find that a mutation that decreases the level of FACT (*pob3-L78R*) results in increased levels of H2B-Ub that is comparable to a loss of Ubp10 (Figure 4). We also show that combining FACT mutations with loss of Ubp10, but not loss of Ubp8, causes sensitivity to hydroxyurea, suggesting a role for Ubp10 in DNA replication. Similarly, combining FACT mutations with deletion of *UBP10*, but not *UBP8*, activate a cryptic promoter reporter gene, reflecting a role for Ubp10 in maintaining nucleosome occupancy during transcription. These combined *in vivo* and *in vitro* observations point to a role for Ubp10 and FACT in jointly maintaining chromatin organization and have important implications for the different cellular roles of the two H2B-Ub DUBs, Ubp8 and Ubp10.

Our results are consistent with a model in which Ubp10 plays a global role in regulating nucleosome dynamics in concert with FACT, while Ubp8 plays a more restricted role at sites of transcription initiation. This view is consistent with previous observations that Ubp8 deletion leads to higher levels of H2B-Ub in the vicinity of the +1 nucleosome, whereas Ubp10 deletions exhibit broader enrichment of H2B-Ub in gene bodies, particularly in longer open reading frames (Schulze et al., 2011). Ubp8 is targeted to genes in the context of the SAGA complex, a global transcription factor that associates with promoter regions of virtually all RNA polymerase II genes (Baptista et al., 2017, 2018; Bonnet et al., 2014; Venters et al., 2011), which accounts for the observed pattern of H2B-Ub enrichment in ubp8 deletion strains (Schulze et al., 2011). The genome-wide pattern of H2B-Ub enrichment found in *ubp10* deletion strains (Schulze et al., 2011) can be explained by a global function for Ubp10 in nucleosome assembly through its partnership with FACT. FACT has been implicated in transcription through its increased association with frequently transcribed genic regions (Feng et al., 2016; Mayer et al., 2010; Pathak et al., 2018; Saunders et al., 2003), but these same studies show that it is also abundant in intergenic regions. FACT physically interacts with both the MCM replicative helicase and with DNA polymerase *α*, and it promotes rapid deposition of nucleosomes in an *in vitro* replication system (Wittmeyer and Formosa, 1997; Yang et al., 2016; Yeeles et al., 2017). FACT is found in yeast cells at about two-thirds the abundance of nucleosomes and therefore is a global component of chromatin that participates in a broad range of chromatin-dependent processes (Gurova et al., 2018). These roles include restoring chromatin integrity after disruptions like transcription, but emerging evidence shows that FACT may have a less prominent role in proliferation of differentiated mammalian cells (Gurova et al., 2018). Collectively, these findings suggest, instead, that the primary role of FACT is to stabilize or maintain existing global chromatin architectures and promote transitions to new patterns during differentiation. The finding that Ubp10, but not Ubp8, plays an unanticipated role in DNA replication (Figures 4 and 5) and suppression of cryptic transcription (Figure 6) that overlaps with the role of FACT supports the idea that, like FACT, Ubp10 has a global function in maintaining stable chromatin architecture against a range of potential perturbations.

In addition to its role in regulating global levels of H2B-Ub, Ubp10 regulates spreading of heterochromatic silencing at telomeres and at mating type loci (Emre et al., 2005; Gardner et al., 2005). This function is mediated by the N-terminus of Ubp10, which is recruited by the SIR complex to subtelomeric regions (Emre et al., 2005; Gardner et al., 2005; Zukowski et al., 2018). The silencing and global H2B ubiquitination functions of Ubp10 appear to be separable, as expression of N-terminal truncations of Ubp10 can restore wild type levels of H2B ubiquitination but are still defective in regulation of subtelomeric silencing (Reed et al., 2015). A recent study (Zukowski et al., 2018) reported that Sir2/Sir4, a subset of the SIR complex, stimulates the activity of GST-Ubp10 on both human nucleosomes and H2A/H2B-Ub heterodimers containing a hydrolyzable non-native linkage between ubiquitin and H2B. The observed stimulation was attributed to the ability of Sir2/Sir4 to recruit Ubp10 to chromatin, as has also been observed in vivo (Emre et al., 2005), as well as to a proposed allosteric stimulation of Ubp10 on both H2A/H2B-Ub heterodimers and on ubiquitinated nucleosomes (Zukowski et al., 2018). The effect observed in that study depended on residues 109 – 133 of Ubp10, which do not play a role in either Ubp10 catalytic activity on yeast H2A/H2B-Ub heterodimers or the ability to be stimulated by FACT (Figure 3). The effect of Sir2/Sir4 on Ubp10 thus appears to be restricted to its silencing-specific functions. Notably, human Usp15 also preferentially deubiquitinates H2A/H2B-Ub dimers and is stimulated by the splicing factor SART3 (Long et al., 2014), although in this case SART3 enhances the activity of Usp15 on H2A/H2B-Ub but does not impact its ability to act on nucleosomes.

It is not clear why nucleosomes are such a poor substrate for Ubp10 as compared to H2A/H2B-Ub heterodimers. Ubp8 is targeted to nucleosomes in the context of the SAGA DUB module, in which Ubp8 forms a complex with Sgf11, Sus1 and Sgf73 (Henry et al., 2003; Kohler et al., 2010). The specificity of Ubp8 for H2B-K123 is conferred by Sgf11, which has a zinc finger domain with an arginine-rich patch (Kohler et al., 2010; Samara et al., 2010a) that docks in the nucleosome acidic patch between histones H2A and H2B (Morgan et al., 2016). By contrast, Ubp10 is a monomeric enzyme that lacks accessory proteins that could promote binding to nucleosomes. However, Ubp10 has surprisingly high affinity for nucleosomes, binding with an apparent Kd of ~400 nM (Figure 3B). The wild-type Ubp8/DUB module, by contrast, does not bind detectably to nucleosomes by EMSA (Figure 3 – Figure supplement 3). The ability of Ubp10 to bind nucleosomes and the stimulation of Ubp10 enzymatic activity by FACT do not seem to be connected, as N-terminal deletions of Ubp10 that impair nucleosome binding are still stimulated by FACT (Figures 3B-D). Moreover, deletions within the N-terminal unstructured region of Ubp10 (Reed et al., 2015), which include residues important for recruitment to telomeres by the Sir complex (Kahana and Gottschling, 1999), have little or no effect on global H2B ubiquitination (Gardner et al., 2005). The relationship between the ability of Ubp10 to bind nucleosomes and use nucleosomes as substrates for deubiquitination therefore remains unresolved.

How does FACT stimulate the activity of Ubp10 on nucleosomes? The structural changes that occur in canonical nucleosomes upon FACT binding are likely key to addressing this question, as the observed stimulation is specific to nucleosomal substrates. FACT can reorganize the nucleosome by disrupting histone/DNA contacts, leaving the nucleosome in an “open” or “destabilized” state (Chen et al., 2018; Kemble et al., 2015; McCullough et al., 2011) and by fully displacing H2A/H2B heterodimers (Belotserkovskaya et al., 2003; Chen et al., 2018; Hsieh et al., 2013; Orphanides et al., 1999; Wang et al., 2018; Xin et al., 2009a). The most straightforward explanation for the effect of FACT on Ubp10 activity is that FACT increases H2B deubiquitination by evicting H2A/H2B-Ub heterodimers, which are an excellent substrate for Ubp10 (Figures 2A and 3C). However, H2A-H2B displacement has been estimated to be limited to 20-50% of the total heterodimers (Orphanides et al., 1998; Xin et al., 2009a) whereas we observe complete deubiquitination, suggesting that dimer displacement can explain some, but not all of the effect of FACT. While it is not known whether ejected H2A/H2B heterodimers retain the H2B-K123 ubiquitin modification *in vivo*, our in vitro results clearly show that an H2A/H2B-Ub dimer bound to FACT can be rapidly deubiquitinated by Ubp10. Alternatively, FACT may stimulate Ubp10 by partly unwinding the DNA (Kemble et al., 2015; McCullough et al., 2011; Valieva et al., 2016), generating a hexasome, or yielding some other intermediate in nucleosome disassembly that either exposes surfaces that allow Ubp10 to interact more favorably with the H2B-Ub linkage or relieves steric clashes. In this second scenario, the assumption is still that Ubp10 deubiquitinates H2B only after FACT has begun to disassemble the nucleosome. In either scenario, Ubp10 must deubiquitinate H2B-Ub at some point between the time the nucleosome is disassembled and before it is reassembled in the wake of either DNA or RNA polymerase. We think it is unlikely that FACT stimulates Ubp10 by recruiting it to the nucleosome, as we see no evidence that FACT enhances Ubp10 binding (Figures 3B and Figure 3 – Figure supplement 2). The enhancement of Ubp10 activity provides the first example of a physiologically relevant substrate that appears to be activated by FACT-mediated reorganization of nucleosomes. Further studies will be needed to unravel the molecular determinants governing Ubp10 substrate preference as well as the mechanism by which FACT activates deubiquitination of nucleosomes.

The coupling of Ubp10 and FACT activity provides a missing link between cycles of H2B ubiquitination and deubiquitination, FACT activity and nucleosome disassembly and assembly that allows us to propose a global model for the role of H2B ubiquitination in chromatin dynamics (Figure 7). Ubiquitination of H2B by Rad6/Bre1 recruits FACT, which facilitates nucleosome disassembly. Ubp10 then deubiquitinates H2B, enhancing the ability of FACT to promote reassembly of the nucleosome and/or reinsertion of a deubiquitinated H2A/H2B dimer. Deubiquitination of H2B could potentially favor nucleosome reassembly by enhancing release of FACT from its proposed checkpoint function (McCullough et al., 2018). The sequence of H2B ubiquitination, FACT-mediated nucleosome reorganization, then deubiquitination by Ubp10 and accelerated assembly could propel sequential assembly of nucleosomes, either in the wake of RNA polymerase or during DNA replication. This proposed sequence of events could also occur in humans, as well, given the conservation of all components of this sytem: RAD6B/RNF20/40 are the human E2/E3, USP36 is the homologue of Ubp10 and human FACT, Spt16/SSRP1, is the homologue of the yeast FACT complex, Spt16/Pob3/Nhp6. Our study provides a framework for understanding how H2B-Ub deubiquitination is coupled to the activity of the histone chaperone, FACT, in producing dynamic changes to nucleosome structure, and has exciting implications for understanding the mechanism by which dynamic cycles of ubiquitination and deubiquitination regulate chromatin organization.

**Figure 7:**
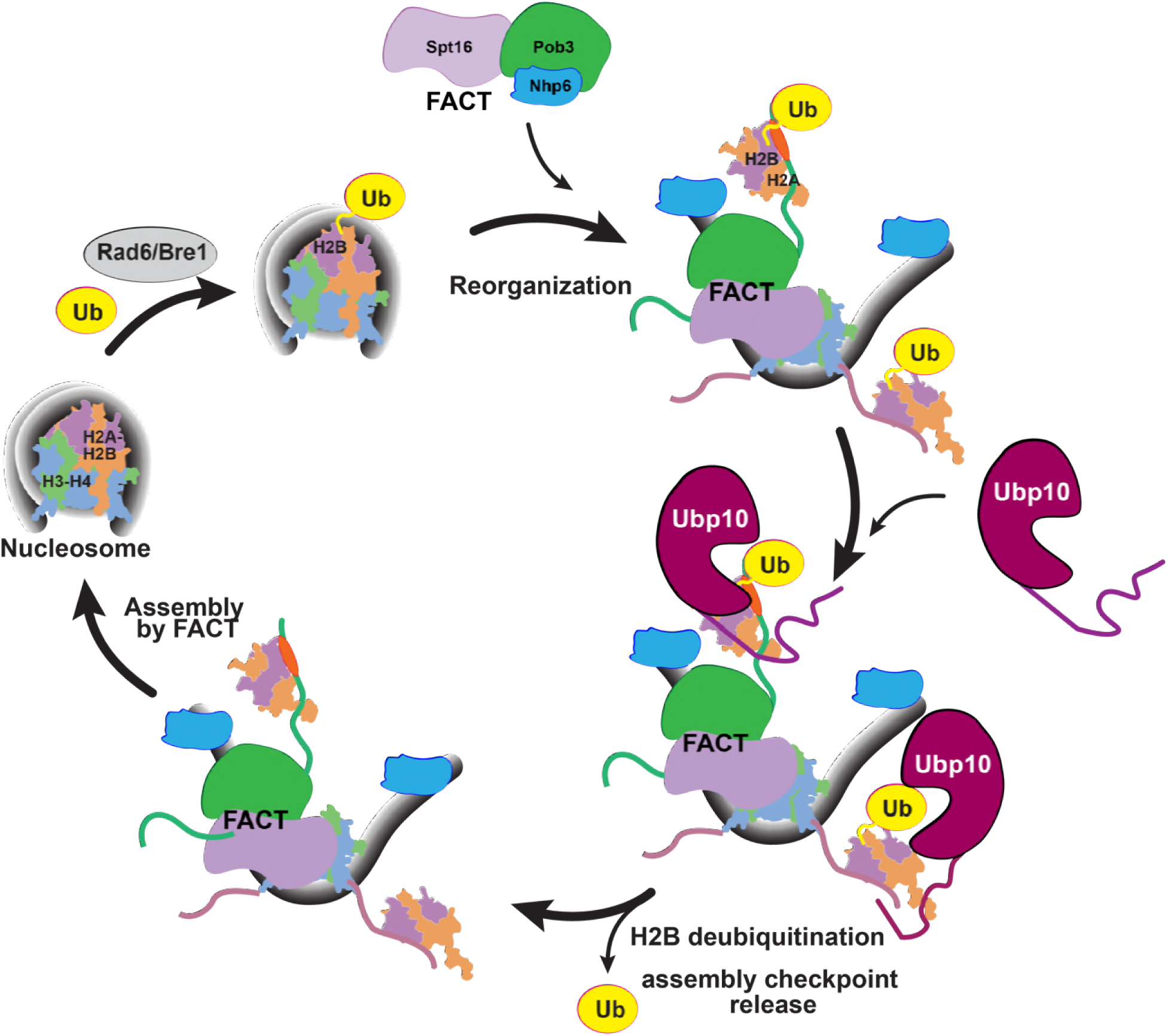
Model for FACT/Ubp10 coordinated H2B deubiquitination and nucleosome assembly during transcription and DNA replication. Rad6/Bre1 ubiquitinates nucleosomal histone H2B during transcription and DNA replication. The presence of H2B-Ub recruits FACT to the nucleosome. FACT reorganizes the nucleosome, displacing some H2A/H2B-Ub heterodimers. Ubp10 deubiquitinates H2B-Ub in the context of fully or partially evicted H2A/H2B-Ub heterodimers and the reorganized nucleosomes. Deubiquitination of H2B signals passage of polymerases and deposition of histones in the wake of polymerases. The deubiquitinated nucleosome is reassembled by FACT, followed by dissociation of FACT.

## Materials and Methods

### Protein expression and purification

#### Purification of Ubp10-WT and Ubp10 truncations

To make the full-length wild-type Ubp10 expression plasmid (pMN2), the protein coding sequences were amplified from *Saccharomyces cerevisiae* genomic DNA by PCR using KOD polymerase (EMD Millipore). The amplified product containing an N-terminal His6-tag and TEV (tobacco etch virus) cleavage site was inserted into a vector that contains thioredoxin protein, pET32a, using IN-fusion cloning kit (Clontech). Ubp10 N-terminal deletions containing residues 157-792 (pMN3), 200-792 (pMN4), and 250-792 (pMN5), were similarly amplified from the original Ubp10-WT expression plasmid, pMN2, and inserted into pET32a. Ubp10-containing plasmids were expressed in Rosetta (DE3) cells. Briefly, a starter culture was grown to an OD of 0.6, then transferred to 1 L M9ZB media and allowed to grow at 37 °C. When the OD reached ~1.5-2, the media was supplemented with 1 mM isopropyl-*β*-D-thiogalactoside (ITPG) and the temperature was shifted to 20°C for an overnight induction. Pelleted cells were lysed in lysis buffer, 25 mM HEPES pH 7.5, 20 mM imidazole pH 7.5, 600 mM NaCl, 10 mM 2-mercaptoethanol, and 1 mM phenylmethylsulfonyl fluoride (PMSF). The lysate was recovered by centrifugation and the supernatant was loaded onto 5 ml HisTrap HP column (GE Healthcare) using buffer A (25 mM HEPES pH 7.5, 20 mM imidazole pH 7.5, 600 mM NaCl, 10 mM 2-mercaptoethanol). Bound protein was eluted with buffer B (25 mM HEPES pH 7.5, 300 mM imidazole pH 7.5, 600 mM NaCl, 10 mM 2-mercaptoethanol). To cleave the purification tags, 1 mg of TEV protease (per mg of protein) was added to the combined fractions and dialyzed overnight against buffer A. The dialyzed sample was then reloaded onto a HisTrap column to remove the cleaved purification tag. The protein was then diluted with ion exchange binding buffer (25 mM HEPES pH 7.5, 50 mM NaCl, 10 mM 2-mercaptoethanol, loaded onto 5 ml Hitrap SP HP column (GE healthcare), and eluted with elution buffer (25 mM HEPES pH 7.5, 1 M NaCl, 10 mM 2-mercaptoethanol). Final purification was carried out using preparative grade HiLoad Superdex 200 26/600 (GE healthcare) with a buffer containing 25 mM HEPES pH 7.5, 250 mM NaCl, and 10 mM 2-mercaptoethanol. All Ubp10 constructs were purified using this protocol. Small aliquots were flash frozen using liquid nitrogen. Although the enzyme is very robust, we avoided freeze thawing for re-use.

#### Purification of wild-type Histones

*Saccharomyces cerevisiae* histones H2A, H2B, H3, and H4 were expressed in *E.coli* and purified by standard methods (Dyer et al., 2004) with modifications as described previously (Morgan et al., 2016). All wild-type histone expression plasmids were generous gifts from the laboratory of Greg Bowman.

#### Preparation of non-hydrolyzable monoubiquitinated H2B

DUB-resistant monoubiquitinated yH2B containing a dichloroacetone linkage between ubiquitin and H2B-K123 (yH2B-DCA-Ub) was prepared using the approach previously described for *Xenopus* H2B-DCA-Ub (Morgan et al., 2016).

#### Purification of yeast FACT

All *Saccharomyces cerevisiae* FACT subunits, Nhp6 and the heterodimer of Spt16 and Pob3 (or mutants) were purified from yeast as previously described (Paull and Johnson, 1995; Xin et al., 2009b).

### Preparation of cleavable monoubiquitinated histone H2B

Ubiquitinated yeast H2B was generated semi-synthetically according to protocols previously reported for *Xenopus* H2B-Ub (Morgan et al., 2016). In brief, ubiquitin (aa1-76, pMN43) and yH2B (aa1-119, pMN161) were cloned into pTXB1 (Evans et al., 1998; Southworth et al., 1999) by making C-terminal fusions with Mxe GyrA intein for thiol-induced cleavage and a chitin binding domain (CBD) for affinity purification using chitin resin (NEB catalog #S6651L). By using this method, both the ubiquitin and H2B carrying a C-terminal reactive thioester are generated, which can then be used in subsequent ligation reactions. We made minor modifications to the previously used purification steps (Morgan et al., 2016). To purify ubiquitin, pMN43 (ubiquitin) was expressed in BL-21 RIL (DE3) cells. A 10-milliliter starter culture was transferred to 1 L 2xYT media, incubated at 37 °C until the OD reached 0.8, and induction was initiated with 0.5 mM IPTG at 16 °C overnight. Pelleted cells were lysed in a buffer containing 100 mM NaOAc, 50 mM HEPES pH 6.5, 0.2 mM PMSF and 1 mM TCEP (Tris(2-carboxymethyl)phosphine hydrochloride). The lysate was spun down and the supernatant was loaded on to chitin resin, incubated for two hours to allow binding, and washed with lysis buffer. MES derivatization was initiated by adding cleavage buffer (100 mM NaOAc, 50 mM HEPES pH 6.5, 0.1 mM PMSF and 0.25 mM TCEP, and 250 mM MESNa (Sodium 2-Mercaptoethanesulfonate) and allowed to proceed at 37 °C overnight. Cleavage was carried out over 5-6 rounds, each time collecting the derivatized ubiquitin by passing it through the chitin resin. The sample was then purified (twice) on an SP column (GE Healthcare), using binding buffer A (50 mM ammonium acetate pH 4.5, 0.5 mM TCEP) and eluted with 10% buffer B (50 mM ammonium acetate pH 4.5, 1 M NaCl, 0.5 mM TCEP). The final sample was thoroughly dialyzed against 0.5% TFA (Trifluoroacetic acid) and lyophilized. Derivatization was verified by MALDI-TOF mass spectrometry.

To purify histone H2B, pMN161 was expressed in BL-21 RIL (DE3) cells. A 10-ml starter culture was prepared from a freshly transformed plate, grown to an OD of 0.4, transferred and inoculated to an OD of 0.6-0.8 in 500 ml 2xYT media, then induced overnight at 25°C by addition of 1 mM IPTG. Cells were pelleted by centrifugation, resuspended in 50 mM Tris-HCl pH 7.5, 200 mM NaCl, 1 mM EDTA, 0.1 mM PMSF, protease inhibitor tablet (1 tablet per 50 ml, Roche), and 0.25 mM TCEP, and lysed with a Microfluidizer (Microfluidics). To prevent degradation, the lysate was immediately spun down, the supernatant was applied on to pre-equilibrated chitin resin, and incubated at room temperature for 2-4 hours. The bound protein was washed with lysis buffer, followed by wash buffer 1 (50 mM Tris-HCl pH 7.2, 200 mM NaCl, 1 mM EDTA, 0.1 mM PMSF and 0.25 mM TCEP), and wash buffer 2 (50 mM Tris-HCl pH 7.4, 200 mM NaCl, 1 mM EDTA, 0.1 mM PMSF and 0.25 mM TCEP). MES derivatization was performed for 18 hours at 4°C in a buffer containing 50 mM Tris-HCl pH 7.4, 200 mM NaCl, 1 mM EDTA, 0.1 mM PMSF, 250 mM MESNa and 0.25 mM TCEP. Cleavage was terminated by adding 20 mM NaOAc pH 5.2, 7 M urea, 1 M NaCl, 1 mM EDTA, 0.1 mM PMSF and 0.25 mM TCEP. The protein was then purified by ion-exchange on an SP column (GE Healthcare) with binding buffer (20 mM NaOAc pH 5.2, 7 M urea, 1 mM EDTA, 0.1 mM PMSF and 0.25 mM TCEP) and eluted with a 0 to 1 M NaCl gradient. Following thorough dialysis against water, the protein was lyophilized and then resuspended in unfolding buffer (7 M Guanidinium-HCl, 20 mM Tris-HCl pH 7.5, and 10 mM DTT). The protein was then purified by HPLC on a C4 column (Higgins Analytical, PROTO C4 5 um 250×10mm) equilibrated with 0.1% TFA and eluted with a 0 to 90% acetonitrile gradient elution. The final sample was then checked for derivatization by MALDI-TOF and immediately lyophilized to prevent hydrolysis.

Synthesis of the C-terminal H2B peptide and ligation of the peptide with H2B and Ub was performed as previously described (Morgan et al., 2016) with the following modifications. Briefly, synthetic peptide Cys-H2B(121-130) and purified thioester peptide H2B-(1-119)-MES were ligated using native chemical ligation, followed by unmasking the protected thiolysine with MgCl_2_ and [Pd(Allyl)Cl]_2_ (Jbara et al., 2016). The ligated product was then treated with DTT, purified via HPLC, and immediately lyophilized. Finally, H2B-(1-130) intermediate was ligated with Ub-MES prepared via intein method and the ligation product was subjected to a desulfurization step, which yielded the desired native H2B-Ub.

### Nucleosome reconstitution

#### H2B-Ub containing nucleosomes

Histone octamers and Widom 601 DNA were purified and reconstituted into nucleosomes using standard methods (Dyer et al., 2004). Nucleosomes containing H2B-DCA-Ub (non-hydrolyzable linkage) and H2B-Ub with the native isopeptide linkage were also reconstituted using the same method and purified using DEAE-5PW column (Tosoh Bioscience). Reconstituted nucleosomes were stored at 4 °C and used as needed.

#### Flag-tagged yeast nucleosomes

Flag-tagged histone octamers were purified from *E.Coli* using a polycistronic expression vector containing all four yeast histones (a generous gift from Alwin Köhler) and a purification tag on H2B as previously described (Turco et al., 2015). Nucleosomes were reconstituted and purified using standard methods.

### Electrophoretic mobility shift assays (EMSAs)

Ubiquitinated or wild-type nucleosomes (100 nM) and Ubp10 concentrations ranging from 0-1600 nM were incubated on ice for 1 hour in the presence of 20 mM HEPES pH 7.6, 50 mM NaCl, 5% sucrose, 1 mM DTT, and 2.5 mM MgCl_2_ and 0.1 mg/ml bovine serum albumin (BSA). Once the reaction was completed, the samples were immediately loaded on to a pre-run 6% Novex TBE gels (Life Technologies) and electrophoresed for 75 minutes using 0.25x TBE running buffer at 4 °C. Gels were stained with SYBR gold (Life Technologies) for 20 minutes and imaged using Chemidoc Touch (Bio-Rad). Apparent dissociation constants were estimated from half-maximal Ubp10-nucleosome complexes on native gel.

### Ubp10 deubiquitination activity assays

Deubiquitination activity assays were performed according to a previously described protocol (Morgan et al., 2016). Briefly, 1 µM yNCP-Ub and 2 µM yH2A/H2B-Ub were preincubated in a 30 °C water bath for 30 minutes in DUB assay buffer (50 mM HEPES pH 7.6, 150 mM NaCl, and 5 mM DTT). Isopeptidase activity was initiated by adding 5 nM pre-warmed (5 minutes) Ubp10. Similar concentrations were used for the experiments involving Ubp10 truncations. For experiments involving FACT, 2 µM Nhp6 & 2 µM Spt16/Pob3 were pre-incubated with the substrates. Time-courses were monitored by removing samples at the indicated times and quenching the reactions with 1x-LDS (Bio-Rad). Samples were analyzed on commercial SDS-PAGE gels (NuPAGE & Criterion) stained with SYPRO Ruby and imaged with Chemidoc Touch (Bio-Rad). All experiments were carried out in siliconized low retention tubes (Fisher Scientific Cat. No.02-681-320).

### Yeast growth assays

Yeast strains with the genotypes shown in Table 1 were grown to saturation in rich medium then 10-fold serial dilutions were spotted to agarose plates with the composition described in Figure 5.

### Western blots

Western blots were performed as in (McCullough et al., 2018) using the TCA method of protein extraction. Each gel contained a dilution series of the WT strain extract to establish linearity of response and to determine the concentration of the target protein.

## Acknowledgements

We thank members of the Wolberger lab for advice and feedback. We are grateful to Anthony DiBello for initially cloning Ubp10 into a bacterial expression plasmid. Funding was provided by National Institutes of Health grants GM095822 (C.W.) and GM064649 (T.F.), the Jordan and Irene Tark Academic Chair (A.B.), an Israel Council of Higher Education Fellowship (M.J.), and a National Science Foundation Graduate fellowship (M.N.).

---

**Figure 2 – Figure supplement 1:**
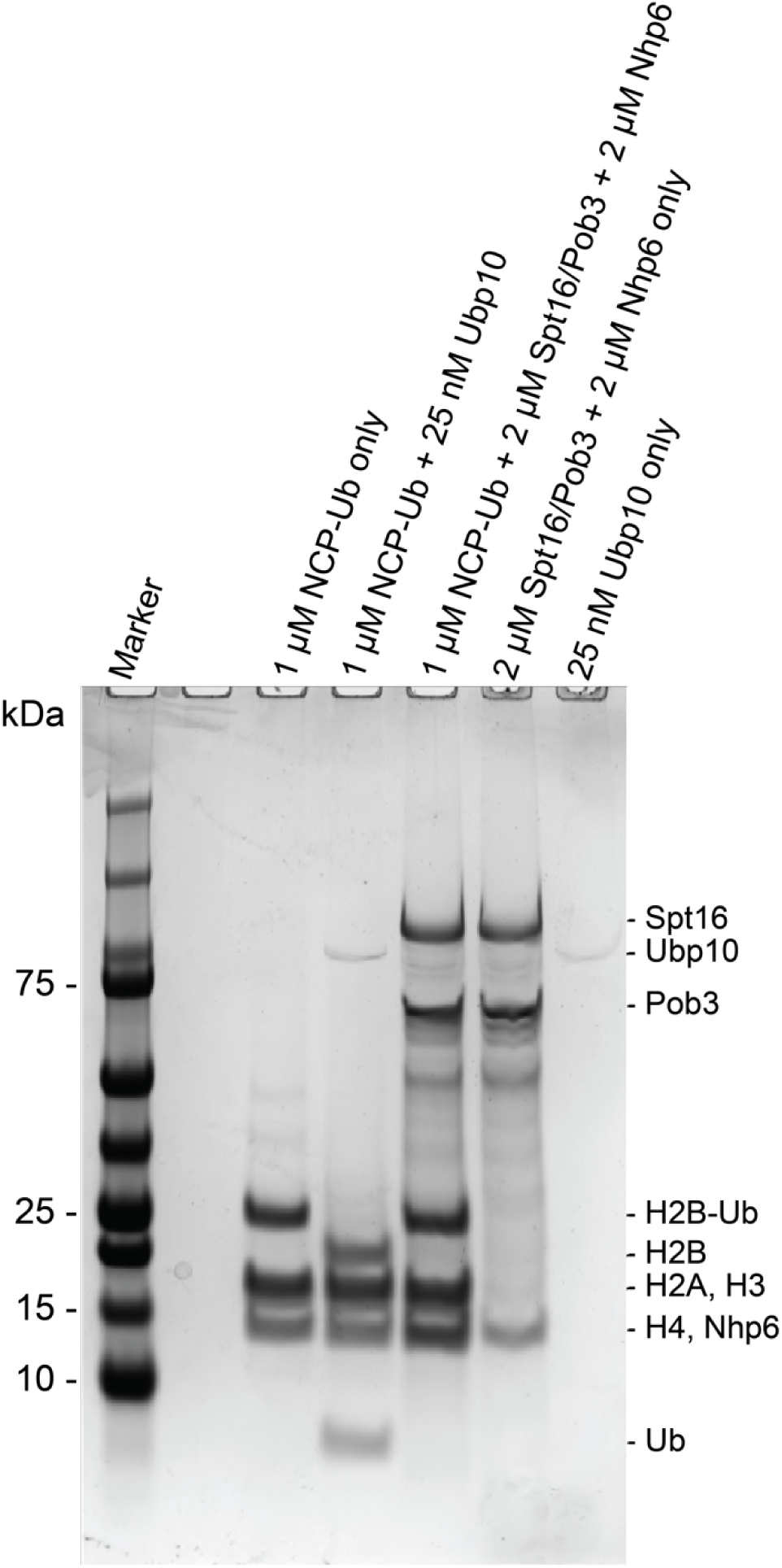
FACT alone does not deubiquitinate H2B-Ub. Control demonstrating that purified Spt16/Pob3 and Nhp6 does not contain contaminants that deubiquitinate H2B-Ub. Proteins indicated in each lane were incubated for 3 hours after which the reactions were quenched. Gel stained with Coomassie.

**Figure 2 – Figure supplement 2:**
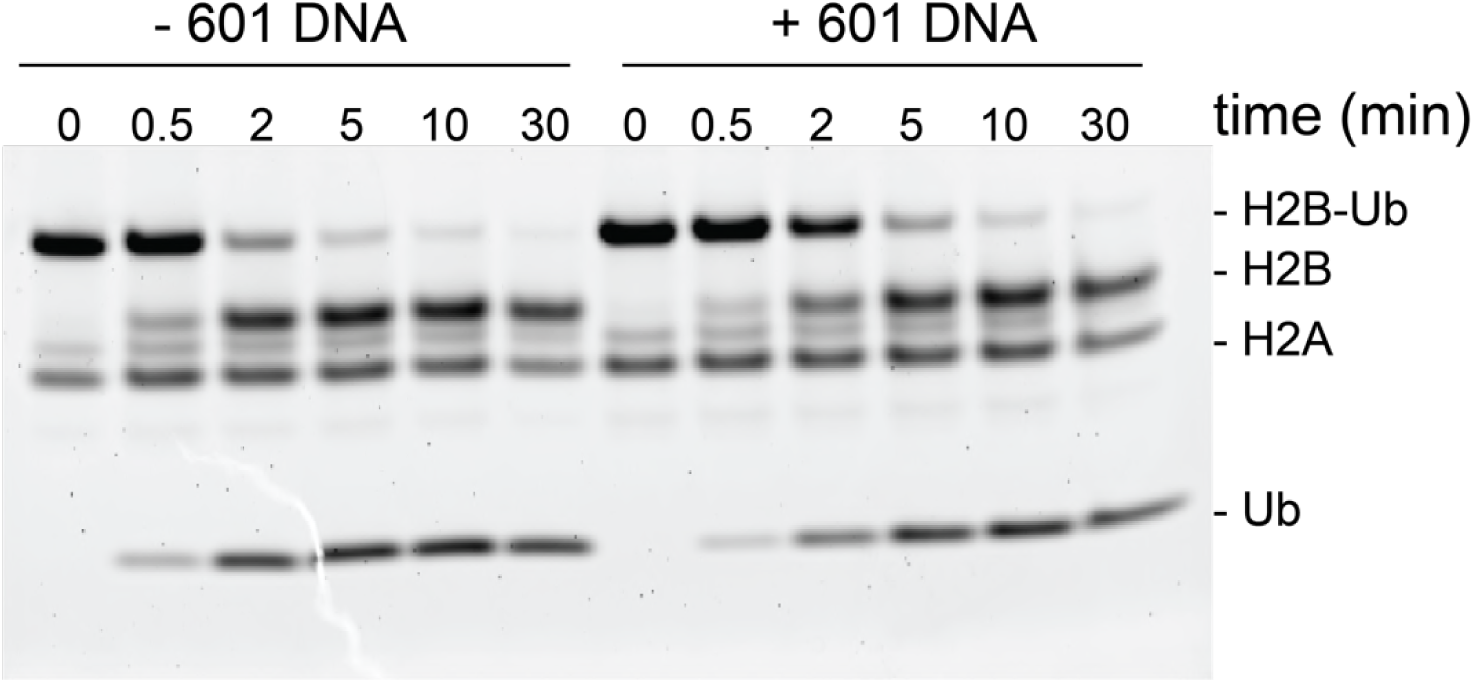
Ubp10 activity on ubiquitinated dimers is unaffected by the presence of 601 DNA. Ubp10 activity on H2A/H2B-Ub with/out 601 DNA. For reaction with 601 DNA, 2 µM Widom 601 DNA was incubated with 2 µM H2A/H2B-Ub for 1 hour. Then the reaction was moved to 30 °C water bath and DUB activity monitored as in Figure 2.

**Figure 3 – Figure supplement 1:**
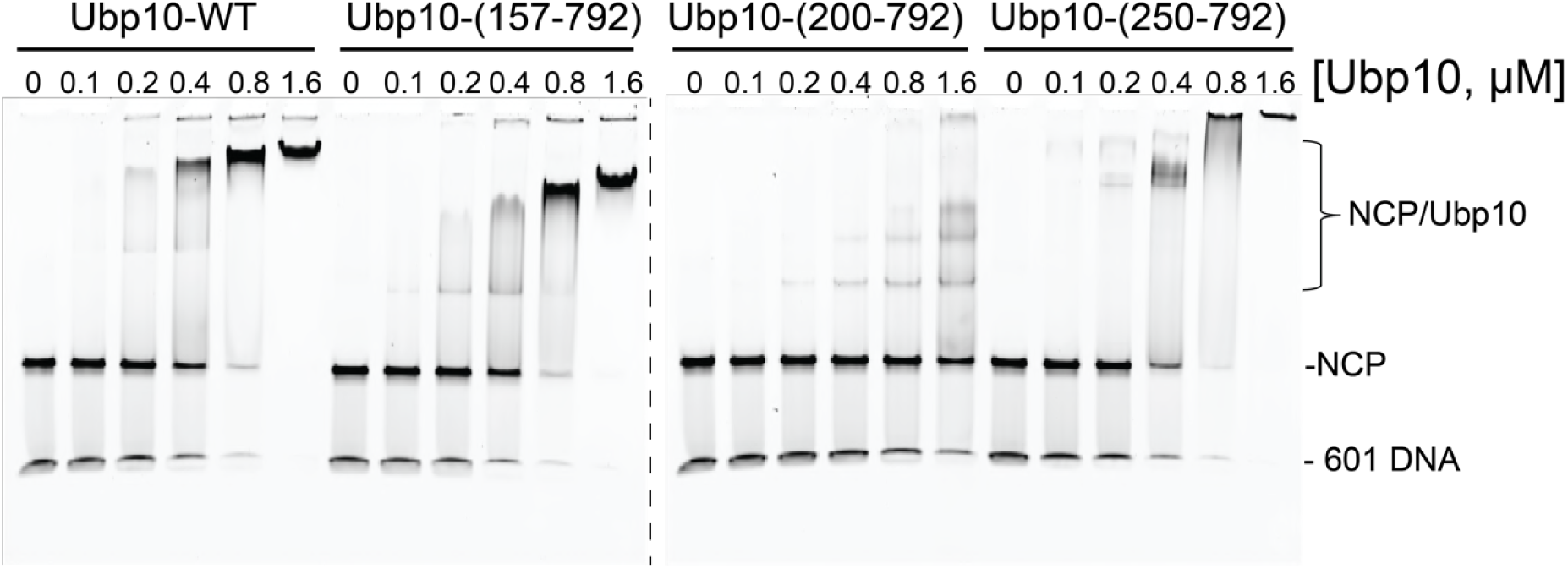
Native gel showing binding of several Ubp10 fragments to unmodified *yeast* nucleosomes. Reaction conditions are the same as in Figure 3B.

**Figure 3 – Figure supplement 2:**
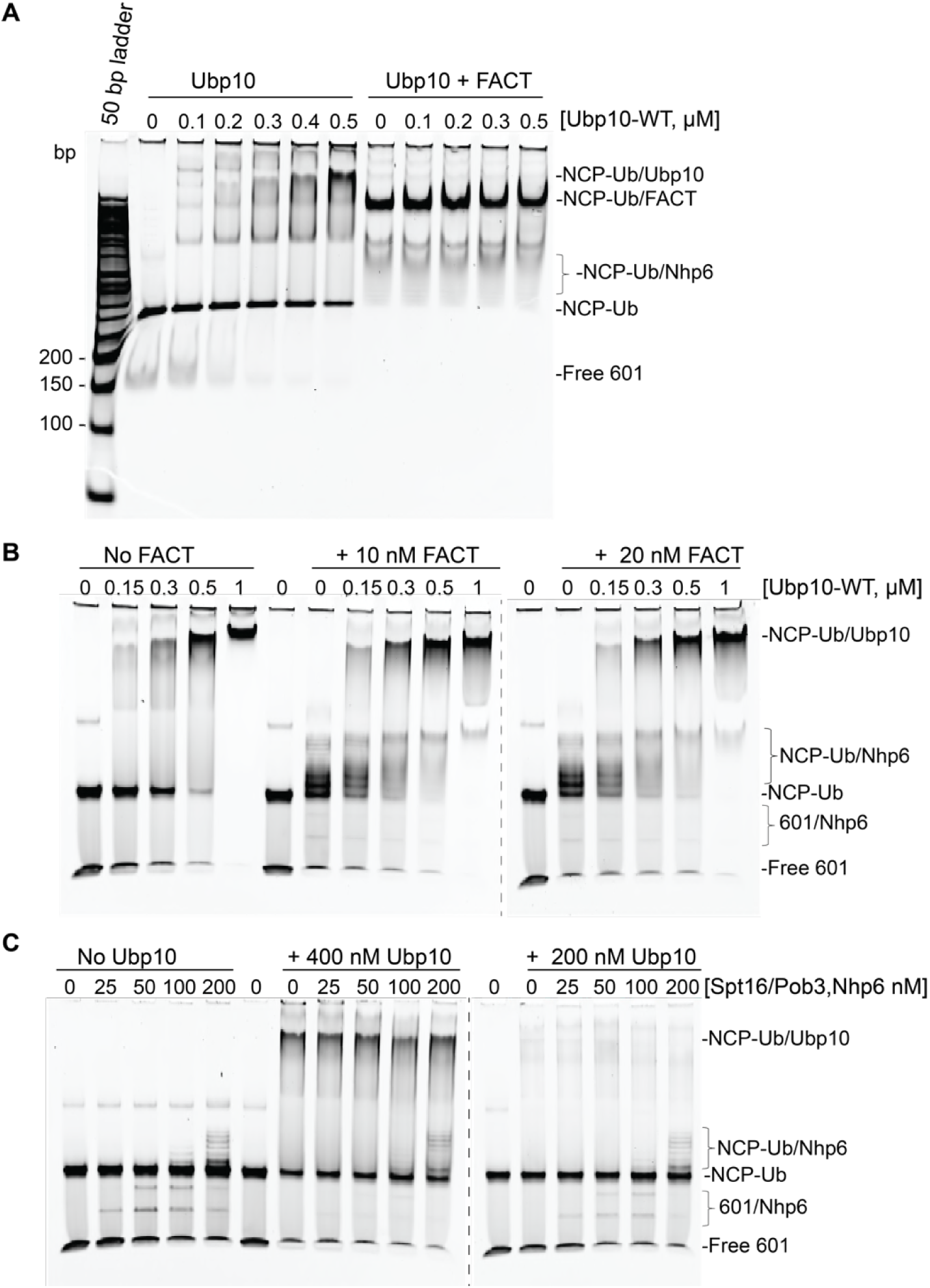
FACT does not affect the affinity of Ubp10 for nucleosomes. EMSA assays of Ubp10 binding to ubiquitinated nucleosomes in the presence and absence of FACT. A) Ubp10/NCP-Ub binding monitored by incubating 100 nM NCP-Ub in EMSA buffer in the absence (left) or in the presence (right) of FACT (2 µM Nhp6 and 2 µM Spt16/Pob3-WT). Samples were incubated for 1 hour on ice and electrophoresed on a 4% acrylamide gel. At high concentration of FACT, the binding of FACT to nucleosomes dominates. B) Reaction in the presence of 2 µM Nhp6 and 10 nM Spt16/Pob3 (left) or 20 nM Spt16/Pob3 (right). C) Reaction with equimolar concentrations of FACT subunits at the apparent K_d_ (left) and below the K_d_ of the binding of Ubp10 to NCP-Ub. Samples in B and C were analyzed on a 6% acrylamide TBE gels (Life Technologies).

**Figure 3 – Figure supplement 3:**
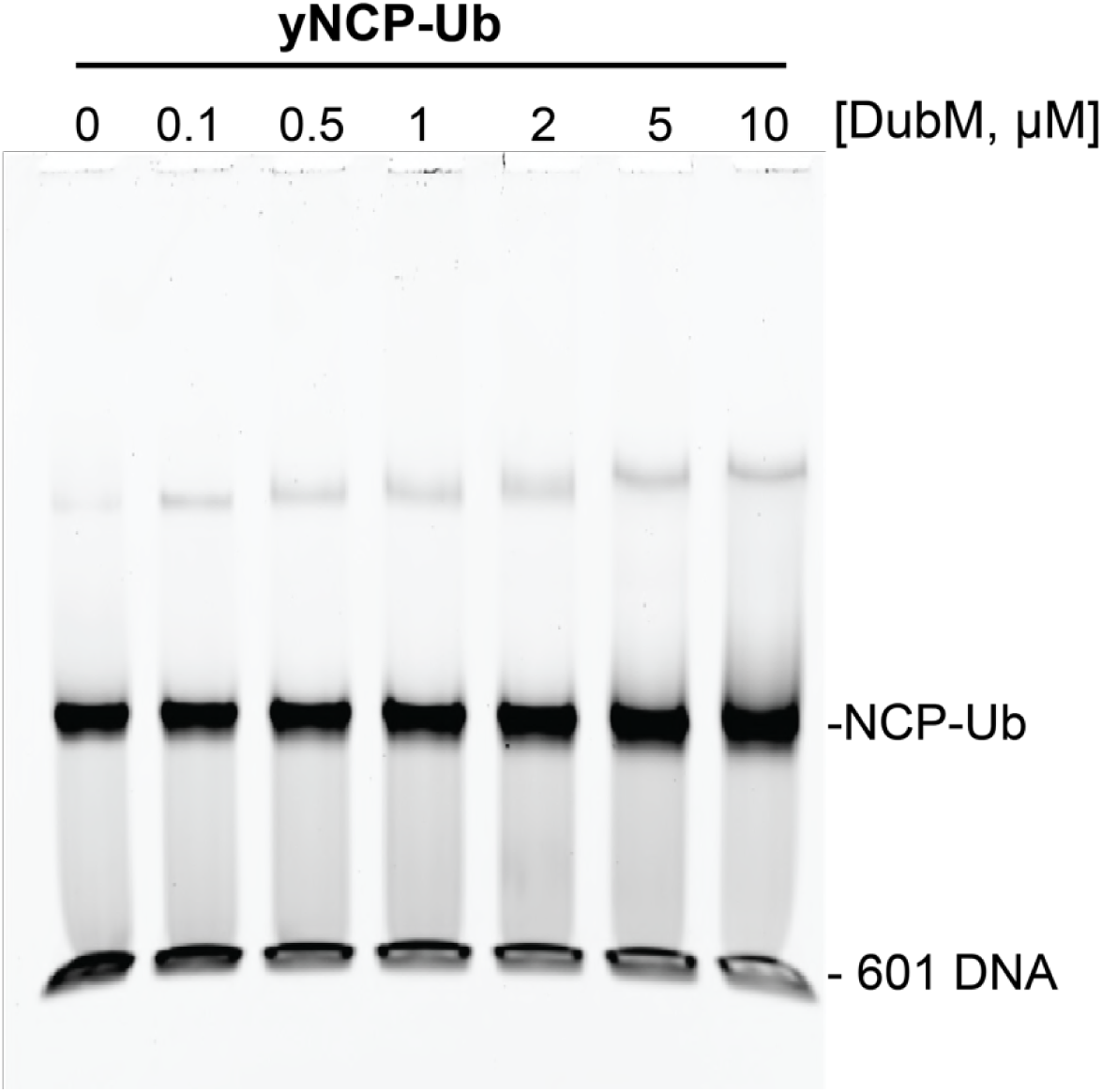
SAGA DUB module binding to ubiquitinated nucleosomes undetectable by EMSA. Increasing amounts of DubM were incubated for 1 hour with 100 nM yeast ubiquitinated nucleosomes (DUB resistant yNCP-DCA-Ub, see methods). Following native gel analysis, the gel was stained with SYBR Gold. The mobility of nucleosomes remained unchaged with increasing concentrations of DUB module.

## References

Bannister, A.J., and Kouzarides, T. (2011). Regulation of chromatin by histone modifications. Cell Res 21, 381–395.

Baptista, T., Grunberg, S., Minoungou, N., Koster, M.J.E., Timmers, H.T.M., Hahn, S., Devys, D., and Tora, L. (2017). SAGA Is a General Cofactor for RNA Polymerase II Transcription. Mol Cell 68, 130–143 e135.

Baptista, T., Grunberg, S., Minoungou, N., Koster, M.J.E., Timmers, H.T.M., Hahn, S., Devys, D., and Tora, L. (2018). SAGA Is a General Cofactor for RNA Polymerase II Transcription. Mol Cell 70, 1163–1164.

Batta, K., Zhang, Z.H., Yen, K.Y., Goffman, D.B., and Pugh, B.F. (2011). Genome-wide function of H2B ubiquitylation in promoter and genic regions. Genes & Development 25, 2254–2265.

Belotserkovskaya, R., Oh, S., Bondarenko, V.A., Orphanides, G., Studitsky, V.M., and Reinberg, D. (2003). FACT facilitates transcription-dependent nucleosome alteration. Science 301, 1090–1093.

Bonnet, J., Wang, C.Y., Baptista, T., Vincent, S.D., Hsiao, W.C., Stierle, M., Kao, C.F., Tora, L., and Devys, D. (2014). The SAGA coactivator complex acts on the whole transcribed genome and is required for RNA polymerase II transcription. Genes Dev 28, 1999–2012.

Bowman, G.D., and Poirier, M.G. (2015). Post-translational modifications of histones that influence nucleosome dynamics. Chem Rev 115, 2274–2295.

Brewster, N.K., Johnston, G.C., and Singer, R.A. (1998). Characterization of the CP complex, an abundant dimer of Cdc68 and Pob3 proteins that regulates yeast transcriptional activation and chromatin repression. J Biol Chem 273, 21972–21979.

Chandrasekharan, M.B., Huang, F., Chen, Y.C., and Sun, Z.W. (2010). Histone H2B C-terminal helix mediates trans-histone H3K4 methylation independent of H2B ubiquitination. Mol Cell Biol 30, 3216–3232.

Chen, P., Dong, L., Hu, M., Wang, Y.Z., Xiao, X., Zhao, Z., Yan, J., Wang, P.Y., Reinberg, D., Li, M., et al. (2018). Functions of FACT in Breaking the Nucleosome and Maintaining Its Integrity at the Single-Nucleosome Level. Mol Cell 71, 284–293 e284.

Cheung, V., Chua, G., Batada, N.N., Landry, C.R., Michnick, S.W., Hughes, T.R., and Winston, F. (2008). Chromatin-and transcription-related factors repress transcription from within coding regions throughout the Saccharomyces cerevisiae genome. PLoS Biol 6, e277.

Cole, A.J., Clifton-Bligh, R., and Marsh, D.J. (2015). Histone H2B monoubiquitination: roles to play in human malignancy. Endocr Relat Cancer 22, T19–33.

Daniel, J.A., Torok, M.S., Sun, Z.W., Schieltz, D., Allis, C.D., Yates, J.R., 3rd, and Grant, P.A. (2004). Deubiquitination of histone H2B by a yeast acetyltransferase complex regulates transcription. J Biol Chem 279, 1867–1871.

Dover, J., Schneider, J., Tawiah-Boateng, M.A., Wood, A., Dean, K., Johnston, M., and Shilatifard, A. (2002). Methylation of histone H3 by COMPASS requires ubiquitination of histone H2B by Rad6. J Biol Chem 277, 28368–28371.

Dyer, P.N., Edayathumangalam, R.S., White, C.L., Bao, Y.H., Chakravarthy, S., Muthurajan, U.M., and Luger, K. (2004). Reconstitution of nucleosome core particles from recombinant histones and DNA. Chromatin and Chromatin Remodeling Enzymes, Pt A 375, 23–44.

Elsasser, S.J., and D’Arcy, S. (2013). Towards a mechanism for histone chaperones. Biochim Biophys Acta 1819, 211–221.

Emre, N.C.T., Ingvarsdottir, K., Wyce, A., Wood, A., Krogan, N.J., Henry, K.W., Li, K.Q., Marmorstein, R., Greenblatt, J.F., Shilatifard, A., et al. (2005). Maintenance of low histone ubiquitylation by Ubp10 correlates with telomere-proximal Sir2 association and gene silencing. Molecular Cell 17, 585–594.

Espinosa, J.M. (2008). Histone H2B ubiquitination: the cancer connection. Genes Dev 22, 2743–2749.

Evans, T.C., Jr., Benner, J., and Xu, M.Q. (1998). Semisynthesis of cytotoxic proteins using a modified protein splicing element. Protein Sci 7, 2256–2264.

Feng, J., Gan, H., Eaton, M.L., Zhou, H., Li, S., Belsky, J.A., MacAlpine, D.M., Zhang, Z., and Li, Q. (2016). Noncoding Transcription Is a Driving Force for Nucleosome Instability in spt16 Mutant Cells. Mol Cell Biol 36, 1856–1867.

Fleming, A.B., Kao, C.F., Hillyer, C., Pikaart, M., and Osley, M.A. (2008). H2B ubiquitylation plays a role in nucleosome dynamics during transcription elongation. Molecular Cell 31, 57–66.

Formosa, T. (2012). The role of FACT in making and breaking nucleosomes. Biochim Biophys Acta 1819, 247–255.

Gardner, R.G., Nelson, Z.W., and Gottschling, D.E. (2005). Ubp10/Dot4p regulates the persistence of ubiquitinated histone H2B: Distinct roles in telomeric silencing and general chromatin. Molecular and cellular biology 25, 6123–6139.

Giannattasio, M., Lazzaro, F., Plevani, P., and Muzi-Falconi, M. (2005). The DNA damage checkpoint response requires histone H2B ubiquitination by Rad6-Bre1 and H3 methylation by Dot1. J Biol Chem 280, 9879–9886.

Gurova, K., Chang, H.W., Valieva, M.E., Sandlesh, P., and Studitsky, V.M. (2018). Structure and function of the histone chaperone FACT - Resolving FACTual issues. Biochim Biophys Acta.

Hahn, M.A., Dickson, K.A., Jackson, S., Clarkson, A., Gill, A.J., and Marsh, D.J. (2012). The tumor suppressor CDC73 interacts with the ring finger proteins RNF20 and RNF40 and is required for the maintenance of histone 2B monoubiquitination. Human Molecular Genetics 21, 559–568.

Henry, K.W., Wyce, A., Lo, W.S., Duggan, L.J., Emre, N.C., Kao, C.F., Pillus, L., Shilatifard, A., Osley, M.A., and Berger, S.L. (2003). Transcriptional activation via sequential histone H2B ubiquitylation and deubiquitylation, mediated by SAGA-associated Ubp8. Genes Dev 17, 2648–2663.

Hsieh, F.K., Kulaeva, O.I., Patel, S.S., Dyer, P.N., Luger, K., Reinberg, D., and Studitsky, V.M. (2013). Histone chaperone FACT action during transcription through chromatin by RNA polymerase II. Proc Natl Acad Sci U S A 110, 7654–7659.

Hwang, W.W., Venkatasubrahmanyam, S., Ianculescu, A.G., Tong, A., Boone, C., and Madhani, H.D. (2003). A conserved RING finger protein required for histone H2B monoubiquitination and cell size control. Mol Cell 11, 261–266.

Jamai, A., Puglisi, A., and Strubin, M. (2009). Histone chaperone spt16 promotes redeposition of the original h3-h4 histones evicted by elongating RNA polymerase. Mol Cell 35, 377–383.

Jasencakova, Z., and Groth, A. (2010). Restoring chromatin after replication: how new and old histone marks come together. Semin Cell Dev Biol 21, 231–237.

Jbara, M., Maity, S.K., Seenaiah, M., and Brik, A. (2016). Palladium Mediated Rapid Deprotection of N-Terminal Cysteine under Native Chemical Ligation Conditions for the Efficient Preparation of Synthetically Challenging Proteins. J Am Chem Soc 138, 5069– 5075.

Jbara, M., Sun, H., Kamnesky, G., and Brik, A. (2018). Chemical chromatin ubiquitylation. Curr Opin Chem Biol 45, 18–26.

Kahana, A., and Gottschling, D.E. (1999). DOT4 links silencing and cell growth in Saccharomyces cerevisiae. Mol Cell Biol 19, 6608–6620.

Kaplan, C.D., Laprade, L., and Winston, F. (2003). Transcription elongation factors repress transcription initiation from cryptic sites. Science 301, 1096–1099.

Kemble, D.J., McCullough, L.L., Whitby, F.G., Formosa, T., and Hill, C.P. (2015). FACT Disrupts Nucleosome Structure by Binding H2A-H2B with Conserved Peptide Motifs. Molecular Cell 60, 294–306.

Kohler, A., Zimmerman, E., Schneider, M., Hurt, E., and Zheng, N. (2010). Structural basis for assembly and activation of the heterotetrameric SAGA histone H2B deubiquitinase module. Cell 141, 606–617.

Komander, D., Clague, M.J., and Urbe, S. (2009). Breaking the chains: structure and function of the deubiquitinases. Nat Rev Mol Cell Biol 10, 550–563.

Kurat, C.F., Yeeles, J.T.P., Patel, H., Early, A., and Diffley, J.F.X. (2017). Chromatin Controls DNA Replication Origin Selection, Lagging-Strand Synthesis, and Replication Fork Rates. Mol Cell 65, 117–130.

Lin, C.Y., Wu, M.Y., Gay, S., Marjavaara, L., Lai, M.S., Hsiao, W.C., Hung, S.H., Tseng, H.Y., Wright, D.E., Wang, C.Y., et al. (2014). H2B Mono-ubiquitylation Facilitates Fork Stalling and Recovery during Replication Stress by Coordinating Rad53 Activation and Chromatin Assembly. Plos Genetics 10.

Long, L., Thelen, J.P., Furgason, M., Haj-Yahya, M., Brik, A., Cheng, D., Peng, J., and Yao, T. (2014). The U4/U6 recycling factor SART3 has histone chaperone activity and associates with USP15 to regulate H2B deubiquitination. J Biol Chem 289, 8916–8930.

Maity, S.K., Jbara, M., and Brik, A. (2016). Chemical and semisynthesis of modified histones. J Pept Sci 22, 252–259.

Malone, E.A., Clark, C.D., Chiang, A., and Winston, F. (1991). Mutations in Spt16/Cdc68 Suppress Cis-Acting and Trans-Acting Mutations That Affect Promoter Function in Saccharomyces-Cerevisiae. Mol Cell Biol 11, 5710–5717.

Mao, P., Meas, R., Dorgan, K.M., and Smerdon, M.J. (2014). UV damage-induced RNA polymerase II stalling stimulates H2B deubiquitylation. Proc Natl Acad Sci U S A 111, 12811–12816.

Mason, P.B., and Struhl, K. (2003). The FACT complex travels with elongating RNA polymerase II and is important for the fidelity of transcriptional initiation in vivo. Mol Cell Biol 23, 8323–8333.

Mayer, A., Lidschreiber, M., Siebert, M., Leike, K., Soding, J., and Cramer, P. (2010). Uniform transitions of the general RNA polymerase II transcription complex. Nat Struct Mol Biol 17, 1272–1278.

McCullough, L., Rawlins, R., Olsen, A., Xin, H., Stillman, D.J., and Formosa, T. (2011). Insight Into the Mechanism of Nucleosome Reorganization From Histone Mutants That Suppress Defects in the FACT Histone Chaperone. Genetics 188, 835–U148.

McCullough, L.L., Connell, Z., Xin, H., Studitsky, V.M., Feofanov, A.V., Valieva, M.E., and Formosa, T. (2018). Functional roles of the DNA-binding HMGB domain in the histone chaperone FACT in nucleosome reorganization. J Biol Chem 293, 6121–6133.

Morgan, M.T., Haj-Yahya, M., Ringel, A.E., Bandi, P., Brik, A., and Wolberger, C. (2016). Structural basis for histone H2B deubiquitination by the SAGA DUB module. Science 351, 725–728.

Moyal, L., Lerenthal, Y., Gana-Weisz, M., Mass, G., So, S., Wang, S.Y., Eppink, B., Chung, Y.M., Shalev, G., Shema, E., et al. (2011). Requirement of ATM-dependent monoubiquitylation of histone H2B for timely repair of DNA double-strand breaks. Mol Cell 41, 529–542.

Ng, H.H., Xu, R.M., Zhang, Y., and Struhl, K. (2002). Ubiquitination of histone H2B by Rad6 is required for efficient Dot1-mediated methylation of histone H3 lysine 79. J Biol Chem 277, 34655–34657.

Orlandi, I., Bettiga, M., Alberghina, L., and Vai, M. (2004). Transcriptional profiling of ubp10 null mutant reveals altered subtelomeric gene expression and insurgence of oxidative stress response. J Biol Chem 279, 6414–6425.

Orphanides, G., LeRoy, G., Chang, C.H., Luse, D.S., and Reinberg, D. (1998). FACT, a factor that facilitates transcript elongation through nucleosomes. Cell 92, 105–116.

Orphanides, G., Wu, W.H., Lane, W.S., Hampsey, M., and Reinberg, D. (1999). The chromatin-specific transcription elongation factor FACT comprises human SPT16 and SSRP1 proteins. Nature 400, 284–288.

Pathak, R., Singh, P., Ananthakrishnan, S., Adamczyk, S., Schimmel, O., and Govind, C.K. (2018). Acetylation-Dependent Recruitment of the FACT Complex and Its Role in Regulating Pol II Occupancy Genome-Wide in Saccharomyces cerevisiae. Genetics 209, 743–756.

Paull, T.T., and Johnson, R.C. (1995). DNA looping by Saccharomyces cerevisiae high mobility group proteins NHP6A/B. Consequences for nucleoprotein complex assembly and chromatin condensation. J Biol Chem 270, 8744–8754.

Pavri, R., Zhu, B., Li, G., Trojer, P., Mandal, S., Shilatifard, A., and Reinberg, D. (2006). Histone H2B monoubiquitination functions cooperatively with FACT to regulate elongation by RNA polymerase II. Cell 125, 703–717.

Ransom, M., Dennehey, B.K., and Tyler, J.K. (2010). Chaperoning histones during DNA replication and repair. Cell 140, 183–195.

Reed, B.J., Locke, M.N., and Gardner, R.G. (2015). A Conserved Deubiquitinating Enzyme Uses Intrinsically Disordered Regions to Scaffold Multiple Protein Interaction Sites. J Biol Chem 290, 20601–20612.

Reinberg, D., and Sims, R.J., 3rd (2006). de FACTo nucleosome dynamics. J Biol Chem 281, 23297–23301.

Robzyk, K., Recht, J., and Osley, M.A. (2000). Rad6-dependent ubiquitination of histone H2B in yeast. Science 287, 501–504.

Ruone, S., Rhoades, A.R., and Formosa, T. (2003a). Multiple Nhp6 molecules are required to recruit Spt16-Pob3 to form yFACT complexes and to reorganize nucleosomes. Journal of Biological Chemistry 278, 45288–45295.

Ruone, S., Rhoades, A.R., and Formosa, T. (2003b). Multiple Nhp6 molecules are required to recruit Spt16-Pob3 to form yFACT complexes and to reorganize nucleosomes. J Biol Chem 278, 45288–45295.

Samara, N.L., Datta, A.B., Berndsen, C.E., Zhang, X., Yao, T., Cohen, R.E., and Wolberger, C. (2010a). Structural insights into the assembly and function of the SAGA deubiquitinating module. Science 328, 1025–1029.

Samara, N.L., Datta, A.B., Berndsen, C.E., Zhang, X.B., Yao, T.T., Cohen, R.E., and Wolberger, C. (2010b). Structural Insights into the Assembly and Function of the SAGA Deubiquitinating Module. Science 328, 1025–1029.

Saunders, A., Werner, J., Andrulis, E.D., Nakayama, T., Hirose, S., Reinberg, D., and Lis, J.T. (2003). Tracking FACT and the RNA polymerase II elongation complex through chromatin in vivo. Science 301, 1094–1096.

Schlesinger, M.B., and Formosa, T. (2000). POB3 is required for both transcription and replication in the yeast Saccharomyces cerevisiae. Genetics 155, 1593–1606.

Schulze, J.M., Hentrich, T., Nakanishi, S., Gupta, A., Emberly, E., Shilatifard, A., and Kobor, M.S. (2011). Splitting the task: Ubp8 and Ubp10 deubiquitinate different cellular pools of H2BK123. Genes & Development 25, 2242–2247.

Simchen, G., Winston, F., Styles, C.A., and Fink, G.R. (1984). Ty-mediated gene expression of the LYS2 and HIS4 genes of Saccharomyces cerevisiae is controlled by the same SPT genes. Proc Natl Acad Sci U S A 81, 2431–2434.

Singh, A., and Xu, Y.J. (2016). The Cell Killing Mechanisms of Hydroxyurea. Genes (Basel) 7, 99.

Southworth, M.W., Amaya, K., Evans, T.C., Xu, M.Q., and Perler, F.B. (1999). Purification of proteins fused to either the amino or carboxy terminus of the Mycobacterium xenopi gyrase A intein. Biotechniques 27, 110–114, 116, 118-120.

Tanny, J.C., Erdjument-Bromage, H., Tempst, P., and Allis, C.D. (2007). Ubiquitylation of histone H2B controls RNA polymerase II transcription elongation independently of histone H3 methylation. Genes Dev 21, 835–847.

Trujillo, K.M., and Osley, M.A. (2012). A role for H2B ubiquitylation in DNA replication. Mol Cell 48, 734–746.

Turco, E., Gallego, L.D., Schneider, M., and Kohler, A. (2015). Monoubiquitination of histone H2B is intrinsic to the Bre1 RING domain-Rad6 interaction and augmented by a second Rad6-binding site on Bre1. J Biol Chem 290, 5298–5310.

Uckelmann, M., and Sixma, T.K. (2017). Histone ubiquitination in the DNA damage response. DNA Repair (Amst) 56, 92–101.

Valieva, M.E., Armeev, G.A., Kudryashova, K.S., Gerasimova, N.S., Shaytan, A.K., Kulaeva, O.I., McCullough, L.L., Formosa, T., Georgiev, P.G., Kirpichnikov, M.P., et al. (2016). Largescale ATP-independent nucleosome unfolding by a histone chaperone. Nat Struct Mol Biol 23, 1111–1116.

VanDemark, A.P., Xin, H., McCullough, L., Rawlins, R., Bentley, S., Heroux, A., Stillman, D.J., Hill, C.P., and Formosa, T. (2008). Structural and functional analysis of the Spt16p N-terminal domain reveals overlapping roles of yFACT subunits. J Biol Chem 283, 5058–5068.

Venters, B.J., Wachi, S., Mavrich, T.N., Andersen, B.E., Jena, P., Sinnamon, A.J., Jain, P., Rolleri, N.S., Jiang, C., Hemeryck-Walsh, C., et al. (2011). A comprehensive genomic binding map of gene and chromatin regulatory proteins in Saccharomyces. Mol Cell 41, 480–492.

Wang, T., Liu, Y., Edwards, G.B., Krzizike, D.D., Scherman, H., and Luger, K. (2018). The histone chaperone FACT modulates nucleosome structure by tethering its components. bioRxiv.

Warfield, L., Ramachandran, S., Baptista, T., Devys, D., Tora, L., and Hahn, S. (2017). Transcription of Nearly All Yeast RNA Polymerase II-Transcribed Genes Is Dependent on Transcription Factor TFIID. Mol Cell 68, 118–129 e115.

Weake, V.M., and Workman, J.L. (2008). Histone ubiquitination: triggering gene activity. Mol Cell 29, 653–663.

West, M.H., and Bonner, W.M. (1980). Histone 2B can be modified by the attachment of ubiquitin. Nucleic Acids Res 8, 4671–4680.

Winkler, D.D., Muthurajan, U.M., Hieb, A.R., and Luger, K. (2011). Histone chaperone FACT coordinates nucleosome interaction through multiple synergistic binding events. J Biol Chem 286, 41883–41892.

Wittmeyer, J., and Formosa, T. (1995). Identifying DNA replication complex components using protein affinity chromatography. Methods Enzymol 262, 415–430.

Wittmeyer, J., and Formosa, T. (1997). The Saccharomyces cerevisiae DNA polymerase alpha catalytic subunit interacts with Cdc68/Spt16 and with Pob3, a protein similar to an HMG1-like protein. Mol Cell Biol 17, 4178–4190.

Wittmeyer, J., Joss, L., and Formosa, T. (1999). Spt16 and Pob3 of Saccharomyces cerevisiae form an essential, abundant heterodimer that is nuclear, chromatin-associated, and copurifies with DNA polymerase alpha. Biochemistry 38, 8961–8971.

Wood, A., Krogan, N.J., Dover, J., Schneider, J., Heidt, J., Boateng, M.A., Dean, K., Golshani, A., Zhang, Y., Greenblatt, J.F., et al. (2003). Bre1, an E3 ubiquitin ligase required for recruitment and substrate selection of Rad6 at a promoter. Mol Cell 11, 267–274.

Xin, H., Takahata, S., Blanksma, M., McCullough, L., Stillman, D.J., and Formosa, T. (2009a). yFACT Induces Global Accessibility of Nucleosomal DNA without H2A-H2B Displacement. Mol Cell 35, 365–376.

Xin, H., Takahata, S., Blanksma, M., McCullough, L., Stillman, D.J., and Formosa, T. (2009b). yFACT induces global accessibility of nucleosomal DNA without H2A-H2B displacement. Mol Cell 35, 365–376.

Yang, J., Zhang, X., Feng, J., Leng, H., Li, S., Xiao, J., Liu, S., Xu, Z., Xu, J., Li, D., et al. (2016). The Histone Chaperone FACT Contributes to DNA Replication-Coupled Nucleosome Assembly. Cell Rep 16, 3414.

Yang, S., Liu, L., Cao, C., Song, N., Wang, Y., Ma, S., Zhang, Q., Yu, N., Ding, X., Yang, F., et al. (2018). USP52 acts as a deubiquitinase and promotes histone chaperone ASF1A stabilization. Nat Commun 9, 1285.

Yeeles, J.T.P., Janska, A., Early, A., and Diffley, J.F.X. (2017). How the Eukaryotic Replisome Achieves Rapid and Efficient DNA Replication. Mol Cell 65, 105–116.

Zhang, X.Y., Varthi, M., Sykes, S.M., Phillips, C., Warzecha, C., Zhu, W., Wyce, A., Thorne, A.W., Berger, S.L., and McMahon, S.B. (2008). The putative cancer stem cell marker USP22 is a subunit of the human SAGA complex required for activated transcription and cell-cycle progression. Mol Cell 29, 102–111.

Zukowski, A., Al-Afaleq, N.O., Duncan, E.D., Yao, T., and Johnson, A.M. (2018). Recruitment and allosteric stimulation of a histone-deubiquitinating enzyme during heterochromatin assembly. J Biol Chem 293, 2498–2509.

